# Connecting single-cell ATP dynamics to overflow metabolism, cell growth and the cell cycle in *Escherichia coli*

**DOI:** 10.1101/2022.05.03.490533

**Authors:** Wei-Hsiang Lin, Christine Jacobs-Wagner

**Affiliations:** Department of Biology, Stanford University, Palo Alto, CA 94305, USA; Chemistry, Engineering, Medicine for Human Health Institute, Stanford University, Palo Alto, CA 94305, USA; Howard Hughes Medical Institute, Stanford University, Palo Alto, CA 94305, USA

## Abstract

Adenosine triphosphate (ATP) is a universal energy-carrying molecule that cells consume and regenerate in vast amounts to support growth. Despite this high turnover, bacterial cultures maintain a similar average concentration of ATP even when the carbon source conditions lead to large differences in population growth rate. What happens in individual bacterial cells is, however, less clear. Here, we use the QUEEN-2m biosensor to quantify ATP dynamics in single *Escherichia coli* cells in relation to their growth rate, metabolism, cell cycle, and cell lineage. We find that ATP dynamics are more complex than expected from population studies and are associated with growth rate variability. Under a nutrient-rich condition, cells can display large fluctuations in ATP level that are partially coordinated with the cell cycle. Abrogation of aerobic acetate fermentation (overflow metabolism) through genetic deletion considerably reduces both the amplitude of ATP level fluctuations and the cell cycle trend. Similarly, growth in media in which acetate fermentation is lower or absent results in reduction of ATP level fluctuation and cell cycle trend. This suggests that overflow metabolism exhibits temporal dynamics, which contributes to fluctuating ATP levels during growth. Remarkably, at the single-cell level, growth rate negatively correlates with the amplitude of ATP fluctuation for each tested condition, linking ATP dynamics to growth rate heterogeneity in clonal populations. Our work highlights the importance of single-cell analysis in studying cellular energetics and its implication to phenotypic diversity and cell growth.

## Introduction

Adenosine triphosphate (ATP) is the most widely used source of cellular energy across all three domains of life. Hydrolysis of ATP, which generates adenosine diphosphate (ADP), inorganic phosphate, and about 30 kJ/mol of free energy, is coupled with a wide range of metabolic processes and cellular functions. Inside cells, ATP can be generated via different processes (e.g., glycolysis, respiration, and fermentation), the choice of which is dictated not only by the metabolic pathways present in the cell, but also by the nature of the carbon source available from the environment. The composition of the carbon source, in turn, has a large impact on cell growth rate. For example, the doubling time of an *Escherichia coli* culture can vary from 20 min to several hours depending on the carbon source^1,2^. Yet, bulk measurements have demonstrated that average cellular concentrations of ATP remain remarkably similar under slow and fast growth conditions^3-5^. Only when cultures run out of an essential nutrient and transition to stationary phase do cellular ATP levels drop by as much as 50 to 80%^6,7^.

In the aforementioned growth studies, the ATP measurements were performed using analytical methods on cell populations. Therefore, they only provide averaged values across millions of cells. What happens at the single-cell level is less clear. An enormous amount of ATP is consumed and synthesized during cell growth. For example, the biomass synthesis occurring during a single cell cycle of *E. coli* is estimated to require about 10^10^ ATP equivalents of energy^8^. Given the millimolar concentration of ATP inside cells^3,9^, the entire pool of ATP molecules in an *E. coli* cell must be consumed and resynthesized over 10,000 times each cell cycle (see Supplementary Information). Such a high turnover raises the question of whether the cellular ATP level might fluctuate during growth in response to changes in cellular demand or metabolic output during the cell cycle. In fact, fluctuations in ATP levels have been reported in eukaryotes. Studies on synchronized cultures have shown that mammalian cells can display oscillatory dynamics of ATP concentration in synchrony with the cell cycle^10^. Oscillatory dynamics have also been reported in synchronized continuous yeast cultures under glucose-limiting condition^11^. When glucose is not limiting, the picture is less clear in yeast as both oscillations and stable concentrations of ATP have been reported at the single-cell level^12,13^. Whether the ATP level oscillates in bacteria is unknown. Dynamic behavior of ATP is not intuitive in these organisms as they lack the complex regulatory systems of cyclins, cyclin-dependent kinases, AMP-activated kinase, and mTOR that connect metabolism to the cell cycle in eukaryotes^14-21^. Interestingly, through the development of a ratiometric fluorescent ATP biosensor, Yaginuma and coworkers^22^ found that the ATP concentration varies widely across cells of an isogenic *E. coli* population. Since the distribution of ATP concentrations was obtained from snapshot microscopy images, it remains to be determined whether this large cell-to-cell variability in cellular ATP levels reflects epigenetic inheritance across cells, random fluctuations, or periodic changes in ATP concentration in individual cells.

The cell-to-cell variability of ATP levels in *E. coli* is further intriguing in the context of cell growth. Under a given growth condition, cells also display considerable variability in growth rate at the single-cell level^23,24^. While single-cell growth rate is correlated between sibling cells and can decrease with cell aging^25-27^, the physiological origin of this cell growth heterogeneity is poorly understood. Whether this cell growth heterogeneity may be connected to the particular ATP concentration or dynamics that each cell experiences has not, to our knowledge, been examined in *E. coli* or any other organism.

Given these many unknowns and the importance of ATP as a key source of cellular energy, we set out to quantify ATP dynamics in single *E. coli* cells in relation to their cell growth rate, metabolism, cell cycle, and cell lineage.

## Results

### ATP levels exhibit large fluctuations during exponential growth

We measured the ATP level inside live *E. coli* cells (strain CJW7548, referred to as “reference” or “ref.” strain hereafter) using QUEEN-2m, a fluorescence biosensor that binds ATP with a *K*_*d*_ ≈ 2 mM^22^. This dissociation constant is well suited for the millimolar ATP level of growing *E. coli* cells^6,9^. QUEEN-2m is a ratiometric biosensor that has excitation peaks at 405 nm and 488 nm corresponding to the ATP-bound and ATP-free forms, respectively. The ratio of fluorescence intensities between the two spectra, denoted as R_ATP_(405ex/488ex) or simply R_ATP_, acts as a proxy for the ATP level and can be converted into absolute ATP concentration after calibration using a luciferase assay (see STAR*Methods and Supplementary Information). We optimized the biosensor expression level and fluorescence exposure times to maximize signal detection over time while minimizing cell phototoxicity and photobleaching (see STAR*Methods). The QUEEN-2m biosensor is sensitive to temperature^22^. Therefore, unless indicated, our experiments were performed at 25°C, which, compared to higher temperatures (such as 37°C), results in stronger fluorescence intensities and greater sensitivity of signal ratio in response to changes in ATP levels^22^. For time-lapse microscopy, we grew cells in the chambers of a custom-made microfluidic device similar to the “mother machine”^24^. In this device, cells were aligned in a single row (Figure 1A), facilitating automated image analysis.

**Figure 1:**
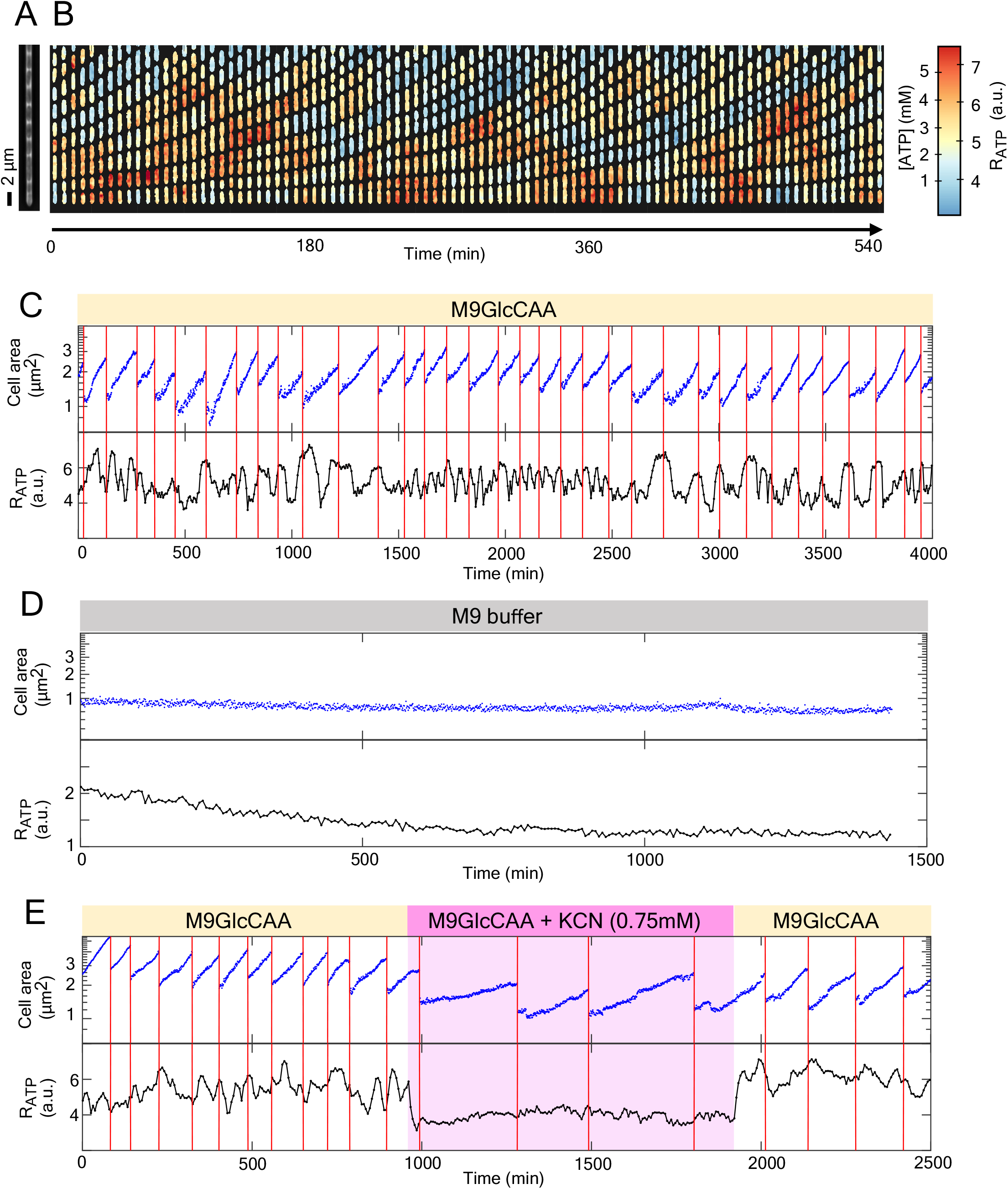
ATP level dynamics in cells under different growth conditions. (A) Phase contrast image of a microfluidic chamber filled with growing *E. coli* cells (CJW7548) at time t = 0 min. (B) Kymograph of ATP dynamics under exponential growth in M9GlcCAA medium. The heatmap shows the scale of R_ATP_ and ATP concentration in each cell over time. (C) Cell size and R_ATP_ dynamics over time along a cell lineage. The red vertical lines indicate the times of cell divisions. (D) Cell size and R_ATP_ dynamics in a stationary phase cell in M9 buffer lacking a carbon source. (E) R_ATP_ dynamics in a cell lineage growing in M9GlcCAA in the absence or presence of KCN. The red vertical lines indicate the times of cell divisions. For panels B-E, a.u. means arbitrary units.

In our first set of experiments, cells were grown in M9 medium containing 0.2% glucose and 0.5% casamino acids (M9GlcCAA), which is a nutrient-rich medium. We found that the R_ATP_ can fluctuate greatly over time (Movie S1), as also shown in kymographs (Figure 1B). This suggests that the ATP level is highly dynamic during exponential growth of *E. coli* in M9GlcCAA. Size measurements of the “mother cell” (located at the terminal position of the microfluidic chamber) over time confirmed the maintenance of exponential growth in the microfluidic chamber (Figure 1C), which was supported by fast nutrient diffusion within the chambers^24^.

Based on our calibration (see Supplementary Information), we estimated that within a cell cycle, the average ATP level is 2.4 mM (ΔR_ATP_ = 4.9 a.u.) and the average peak and valley of the fluctuations are 3.6 mM (R_ATP_ = 5.7 a.u.) and 1.2 mM (R_ATP_ = 4.1 a.u.), respectively. Large fluctuations in ATP concentration persisted for multiple cell cycles over the course of the multi-day microfluidic experiments (Figure 1C).

Given the ratiometric nature of the QUEEN-2m biosensor, the observed temporal ATP fluctuations were unlikely to be caused by a variability in biosensor protein levels. Consistent with this assumption, the levels of ATP and biosensor were not correlated in time (Figure S1). In addition, reducing biosensor protein expression by 75% (by inducing QUEEN-2m protein expression with 15 μM Isopropyl β-D-1-thiogalactopyranoside (IPTG) instead of 60 μM) resulted in a similar magnitude of fluctuation in R_ATP_ (Figure S2A-C). Since the biosensor QUEEN-2m has a weak dependency on pH^22^, we next assessed whether a change in pH may contribute to the observed fluctuation in R_ATP_ using the ratiometric pH biosensor, PHP^28^. We found that the intracellular pH remained largely constant throughout the microfluidic experiment (Figure S3), with a fluctuation of less than ± 0.05 pH unit (see STAR*Methods), consistent with a previous report^22^. Such small pH fluctuations are negligible for the R_ATP_ measurements.

In addition, we ruled out the possibility that small fluctuations in temperature during time-lapse experiments are responsible for the observed changes in R_ATP_ given that the R_ATP_ fluctuations are not synchronized across cells in the microfluidic chambers (Movie S1). We also determined that the R_ATP_ fluctuations are neither caused by measurement noise from the digital camera nor by errors in image analysis (see STAR*Methods). The R_ATP_ fluctuations were also observed in different *E. coli* strain backgrounds (see STAR*Methods and Figure S4). Therefore, we concluded that the observed large fluctuations in R_ATP_ are likely due to bona fide changes of the intracellular ATP level over time, which likely contribute to the large cell-to-cell variability in ATP levels previously reported in cell populations^22^.

### ATP fluctuations are associated with cell growth

Figure 1C shows a typical profile of R_ATP_ levels through multiple rounds of exponential cell growth and divisions in M9GlcCAA. To examine whether the ATP dynamics depend on cell growth, we loaded cells from a stationary-phase liquid culture into the microfluidic device and maintained their non-growing status by flowing M9 buffer (i.e., no carbon source) into the microfluidic device. As expected, the overall R_ATP_ and thus ATP level was significantly lower under these nutrient depletion conditions (Figure 1D), as expected^6^. The fluctuations in R_ATP_ levels were also drastically reduced (Figure 1D), presumably down to measurement noise level. This argues that large ATP fluctuations are associated with cell growth. Consistent with this interpretation, reducing cell growth by exposing cells to a sub-lethal concentration of potassium cyanide (KCN), an inhibitor of the electron transport chain, reduced not only the mean R_ATP_ level but also the amplitude of R_ATP_ fluctuations (Figure 1E).

### The ATP dynamics can be decomposed into distinct oscillatory frequencies

At first glimpse, the ATP dynamics in *E. coli* appeared quite variable over time (Figure 1C). However, upon closer examination, we identified two apparent types of oscillatory behaviors (Figure 2A). The slower oscillation correlated well with the cell cycle period, with peaks in R_ATP_ levels occurring at the same time or close to the time of cell division (Figure 2A). The other oscillatory-like behavior was faster and seemed uncorrelated with the cell cycle (Figure 2A). This visual inspection suggested that the ATP dynamics may result from the superposition of different oscillatory processes.

**Figure 2:**
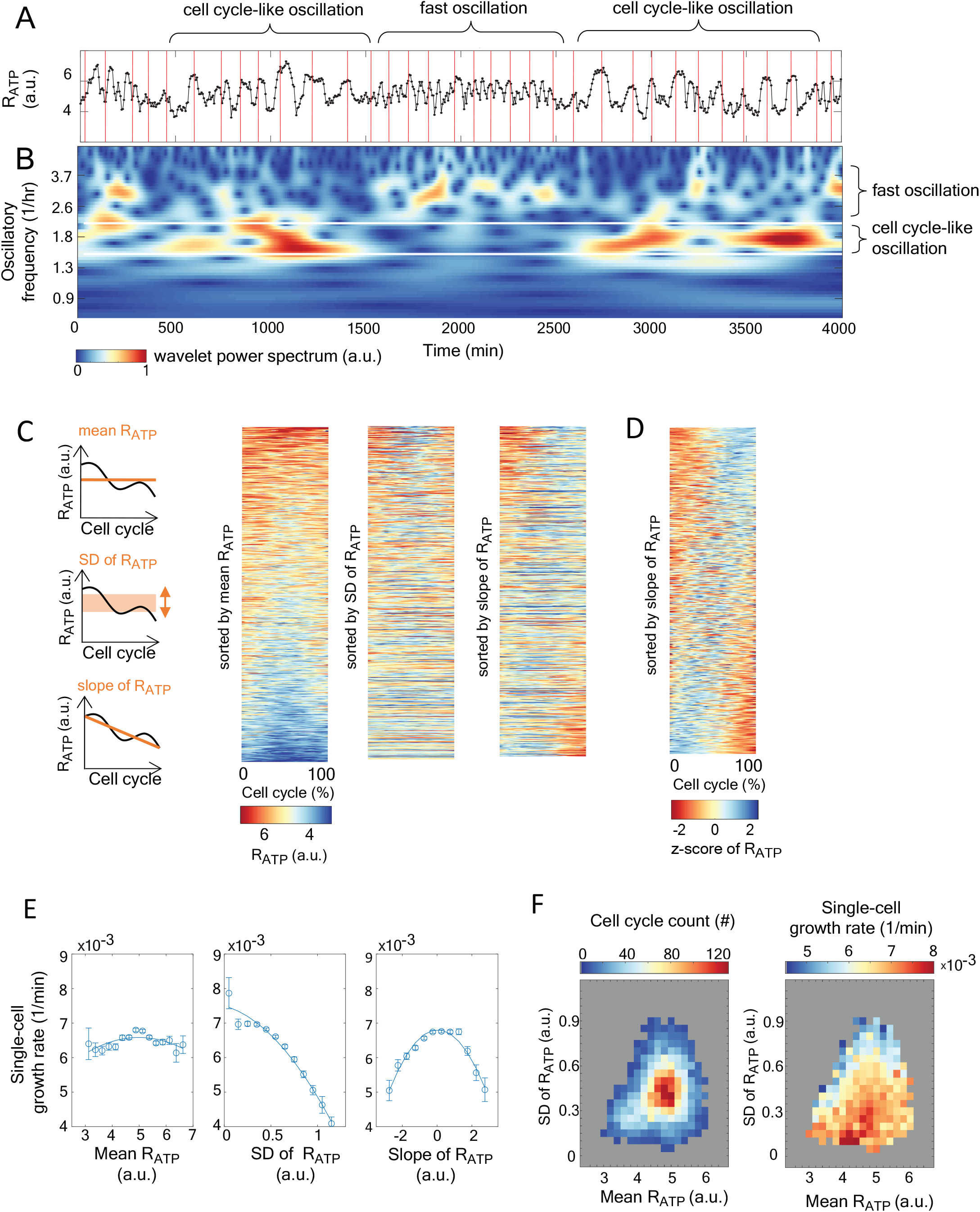
Time-series analysis and cell cycle statistics of ATP and growth rate. (A) A detrended R_ATP_ trajectory (from Figure 1C) of a representative mother cell lineage showing timespans of cell cycle-like oscillations in which R_ATP_ values tend to peak at or near the times of cell division indicated by the red lines. The same trajectory also illustrates a period of faster R_ATP_ oscillations in which more than one R_ATP_ oscillation occurs within a division cycle. (B) Wavelet analysis scalogram of the R_ATP_ time-series shown in (A). Oscillatory signatures are highlighted in the heatmap. (C) Cell-cycle tempograms of absolute R_ATP_ in which each cell cycle pattern was sorted in three different ways, as illustrated in the schematics. Each row in the tempograms (n = 6911 cell cycles) represents one cell cycle (normalized in cell cycle percentile), with the absolute R_ATP_ level shown in the color scale. The rows in left, middle and right tempograms are sorted according to the mean, SD, or the slope of R_ATP_, respectively. (D) Cell-cycle tempogram of the z-transformed R_ATP_ pattern. Same data as in (C) except that z-transformed R_ATP_ is shown in the color scale. (E) Plots showing growth rate as a function of the mean R_ATP_, the SD of R_ATP_, or the slope of the R_ATP_ along the cell cycle (left, middle and right panels, respectively). Data points are binned according to their values in the x-axis. Error bars represent the standard error of the binned data while the solid lines represent the quadratic fits of the binned data. (F) Cell cycle statistics on two-dimensional histograms, where X- and Y-axes depict the mean R_ATP_ and maximal R_ATP_ fluctuation, respectively. The color scale reflects the cell cycle counts (left) and the corresponding single-cell growth rate (right).

To examine the oscillatory dynamics quantitatively, we performed wavelet analysis on individual R_ATP_ dynamics (see STAR* Method). Wavelet analysis detects transient oscillatory patterns in time-series and highlights the signature in a two-dimensional scalogram^29^. Figure 2B illustrates such a scalogram in which the oscillatory signatures are shown for different frequencies (y-axis) and time (x-axis), with the two-dimensional power spectral intensity being shown in a color scale. Consistent with our visual inspection, the wavelet analysis revealed at least two transient oscillatory signals with distinct periods. The slower oscillatory dynamics has a period similar to that of the cell cycle (Figure 2B, white horizontal lines), suggesting a potential cell cycle coupling. The faster dynamics may reflect metabolic oscillations.

To more carefully examine the cell cycle-like oscillatory behavior of R_ATP_ across all recorded cell cycles (n = 6911), we generated three cell-cycle “tempograms” in which each row depicts the R_ATP_ values (in a color scale) along a single cell cycle (0 % = birth and 100% = division) (Figure 2C). These tempograms show the same data sorted based on (i) their mean ATP level (highest to lowest), (ii) magnitude of the ATP level fluctuation represented by the standard deviation (SD) of R_ATP_ values within the cell cycle (highest to lowest) and the trend of the ATP level during assessed by the slope of R_ATP_ through linear regression fitting (descending trend to ascending trend). These tempograms show that the ATP dynamics varied across cell cycles. Some cell cycles displayed overall high R_ATP_ level while others remained low (left tempogram). Some cell cycles exhibited particularly large ATP fluctuations whereas other cell cycles displayed small ones (middle tempogram). In some cases, the ATP level gradually increased or decreased during the cell cycle, while it remained fairly constant in others (right tempogram). Interestingly, when we transformed the R_ATP_ values into z-scores to specifically look at the relative (rather than absolute) levels in ATP in each recorded cell cycle, we found an enrichment in high z-scores of R_ATP_ around either the beginning or the end of the cell cycle (Figure 2D). Thus, regardless of the mean ATP level along the cell cycle, the highest accumulation of ATP tends to occur at or near the time of cell division/birth.

### Growth rate negatively correlates with ATP level fluctuations at the single-cell level

The co-existence of various patterns in absolute ATP levels within an exponentially growing population (Figure 2C) led us to inquire whether the metabolic ATP pattern correlates with the growth rate of the cell. To answer this question, we focused on three ATP statistics across cell cycles (Figure 2E): (i) the mean level of ATP, (ii) SD of R_ATP_ values within the cell cycle, and (iii) the slope of the R_ATP_ data in each cell cycle, as described for the tempograms. For each cell cycle, we compared these ATP statistics to the single-cell growth rates, which we calculated by fitting the cell area over time on a logarithmic scale. We found that the growth rate was not particularly sensitive to the mean ATP level over the cell cycle (Figure 2E, left plot), presumably because the ATP concentration largely remains within a millimolar range that does not impede enzymatic activity and biomass synthesis. In contrast, the growth rate negatively correlated with the degree of ATP fluctuation (SD of R_ATP_) during the cell cycle (Figure 2E, middle plot); on average, the more the ATP level fluctuated within a cell cycle, the slower the cell grew. If this surprising negative correlation was the mere consequence of cell growth being too fast or too slow relative to the biosynthesis rate of ATP, we would expect the ATP concentration to decrease through dilution in cells expanding their volume too quickly. Conversely, we would expect the ATP concentration of cells growing too slowly relative to ATP biosynthesis to increase their ATP concentration through accumulation. However, this is not what is observed (Figure 2E, right plot). The fastest growing cells displayed, on average, a stable ATP level during the cell cycle (i.e., slope close to zero) whereas slow growth rate was associated with both an increase (positive R_ATP_ slope) and decrease (negative R_ATP_ slope) in ATP level. These results may suggest that fluctuations in ATP, regardless of their direction, have a negative impact on growth rate.

Upon further analysis of ATP statistics across cell cycles, we found that, for most cell cycles, the mean ATP level and the ATP fluctuation level are within the intermediate range (Figure 2F, left), as expected for a fairly normal distribution. Interestingly, these prevalent cell cycles tend to be associated with slower growth rates than the less common cell cycles with smaller fluctuations in ATP level (Figure 2F, right). The highest growth rate was associated with the most stable ATP level (i.e., strongest ATP homeostasis) and an intermediate level in ATP.

### Amplitudes of ATP fluctuations are correlated between mother and daughter cells

Lineage analysis of microbial populations has provided evidence of cell aging^26^ and transient memory^30^. To investigate whether the observed ATP statistics are related to lineage structure, we reconstructed cell lineages from our microfluidic records. We found no discernible trend in growth rate, R_ATP_ fluctuations, or mean cell size associated with cell pole age (Figure S5). On the other hand, we found statistical correlations between the behavior of pre-divisional and post-divisional cells (referred to as “mother-daughter pairs”, see Figure 3A) and between post-divisional cells derived from the same cell (referred to as “daughter cell pairs”). Cell growth rate was correlated between mother-daughter pairs and between daughter cell pairs (Figure 3B-C), as previously reported^26,27,31^. The correlation in mean ATP levels between these two pairs was also expected (Figure 3D and E) since metabolites are inherited from mothers to daughters during cell division, resulting in near-equal distribution of cytoplasmic ATP between daughter cells. The correlation of SD of R_ATP_ in mother-daughter and daughter cell pairs was particularly intriguing (Figure 3F and G), since there was no *a priori* reason for the fluctuation of metabolite levels to be inherited. Our data suggest that other unknown variables inherited at division contribute to the correlation in ATP dynamics between mother and daughter cells.

**Figure 3:**
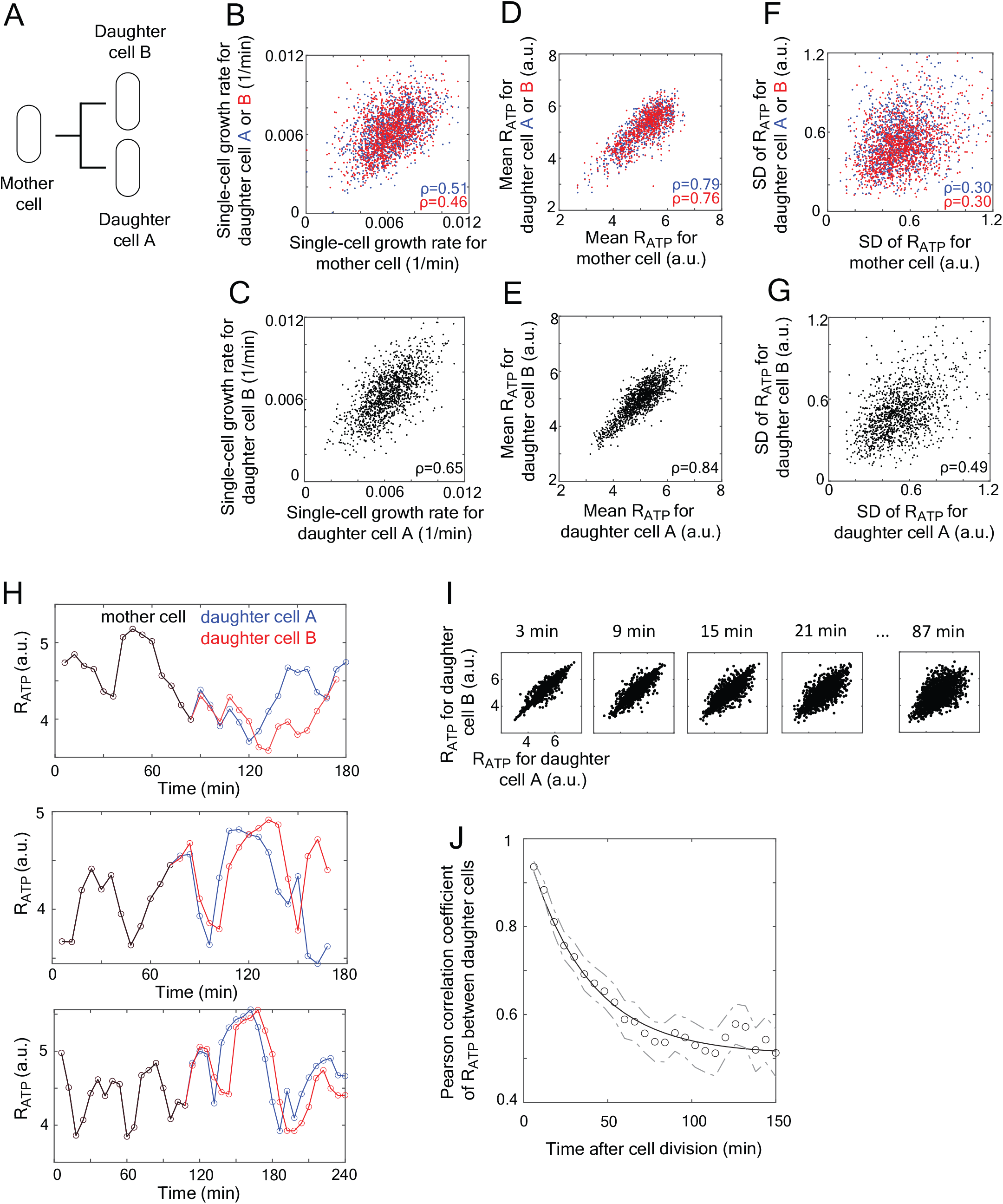
Comparison of ATP dynamics between mother and daughter cells and between daughter cells. The R_ATP_ statistics and dynamics were compared between mother and daughter cells or between daughter cell pairs. (A) Schematic illustrating our definition for a mother cell and its daughter cells A and B. (B-G) Correlation analysis of cell pairs, with ρ representing the Pearson correlation coefficient. (B) Plot showing the single-cell growth rate of mother cells relative to that of their daughter cells A (blue) or B (red) (n = 1306 pairs). (C) Same as in (B) but between daughter cells A and B (n = 1306 daughter cells pairs). (D) Same as in (B) but for the mean of R_ATP_ instead of the single-cell growth rate (E) Same as in (D) but between daughter cells A and B. (F) Same as in (B) but for SD of R_ATP_. (G) Same as in (F) but between daughter cells A and B. (H) Three representative dynamics of R_ATP_ level in a mother cell (black) and its two daughter cells (blue and green). (I) Correlation between R_ATP_ levels of daughter cells pairs at different times after cell division. (J) Decorrelation of R_ATP_ levels between daughter cells pairs over time following cell division. Open circles show the Pearson correlation coefficient of R_ATP_ between daughter cells pairs at a given time while the line represents the fitting of an exponential decay *f*(*t*) = *a* + *be*^*-ct*^. The upper and lower dashed lines indicate the 95% of confidence interval using bootstrapping (without replacement).

### ATP dynamics are correlated between daughter cell pairs

Based on our wavelet analysis (Figure 2B), the intracellular concentration of ATP exhibits transient oscillatory patterns. To determine whether some form of noise caused the observed ATP fluctuations, we compared ATP dynamics within pairs of daughter cells. After cell division, daughter cells become physically separated and the noise component of the ATP fluctuations should become independent and thus uncorrelated. We found that the fluctuation in R_ATP_ levels often remained similar between daughter cell pairs after birth (Figure 3H), with the Pearson’s correlation coefficient (ρ) gradually decaying as the two daughter cells grew independently (n = 1306, Figure 3I). In fact, the correlation remained above 0.5 even ∼100 min after birth (Figure 3J), which corresponds to the end of an average cell cycle under our experimental growth conditions. Since noise cannot be correlated between daughter cell pairs, this result implies that most of the ATP fluctuations derive from physiological regulations rather than cellular or measurement noise.

### The amplitude of ATP fluctuations changes with the quality of the carbon source

We showed that ATP fluctuations dramatically decrease when the mean ATP level and cell growth are reduced, as in starved cells or after metabolic poisoning (Figure 1D and E). This raised the question of whether the amplitude in ATP fluctuations is linked to the growth rate and/or the mean ATP concentration. Previous population measurements have shown that while the type of carbon source can drastically affect the growth rate of *E. coli*, the (mean) ATP level remains similar at the population level^4,5^. Therefore, to examine the potential origin of the ATP fluctuations, we performed additional microfluidic experiments and recorded the R_ATP_ over time in cells growing in M9 media containing different carbon sources, including glucose (M9Glc), xylose (M9Xyl), or glycerol (M9Gly), instead of M9GlcCAA (Figure 4A). Growth rate was, as expected, considerably reduced in M9Glu, M9Xyl and M9Gly compared to the nutrient-richer medium M9GlcCAA (Figure 4B). We also found that, consistent with the literature^4,5^, the mean ATP level remains similar across these growth conditions (Figure 4C). In contrast, the ATP fluctuations were markedly reduced in cell lineages growing in M9Glc, M9Xyl, or M9Gly compared to M9GlcCAA (Figure 4A), with differences in SD of R_ATP_ of about 40% (Figure 4D). We obtained similar results if we compared the coefficient of variation (CV) of the calibrated ATP concentration instead of the SD of R_ATP_ (Figure S6A).

**Figure 4:**
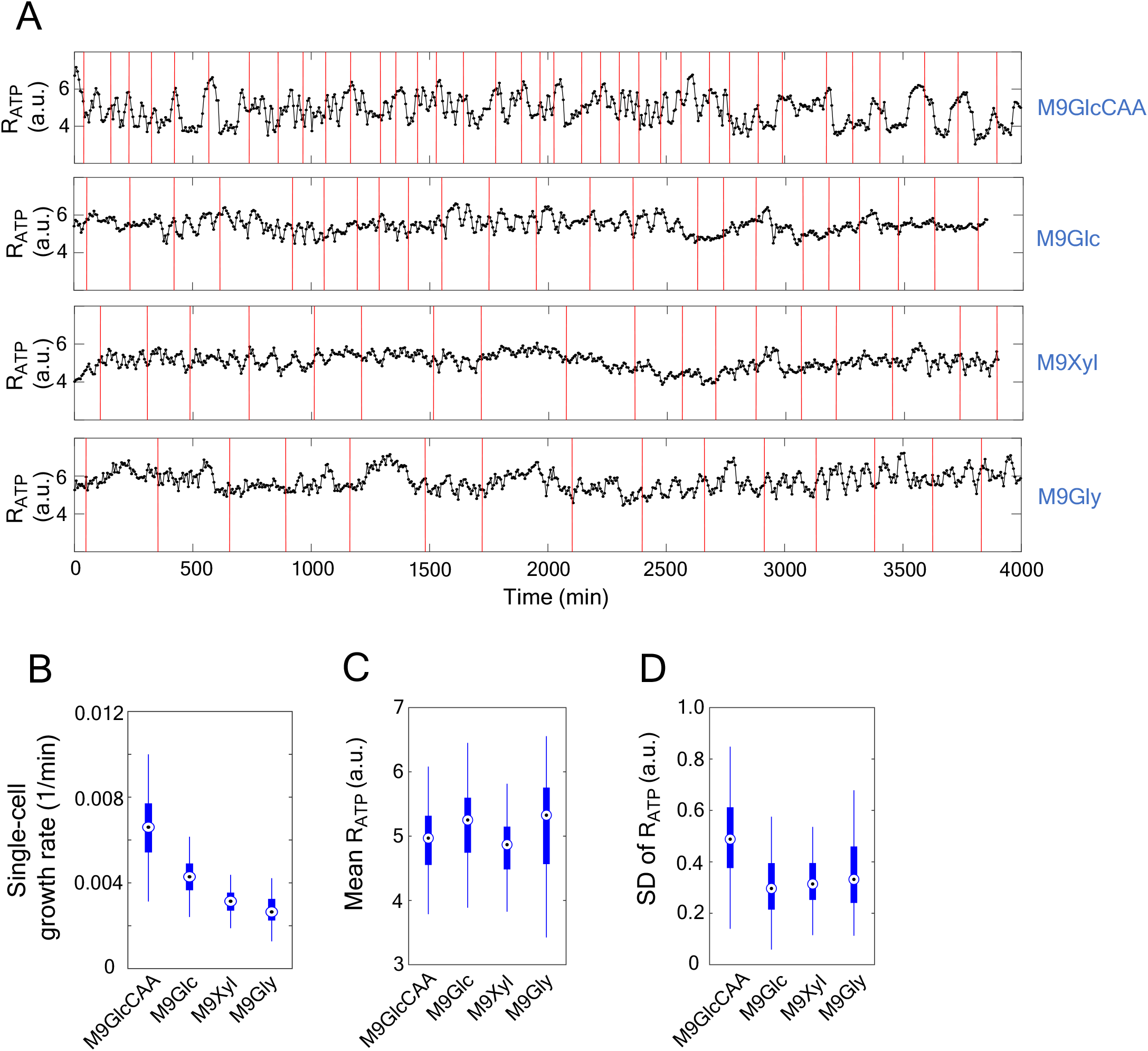
ATP and cell growth dynamics under different nutrient conditions. (A) Representative R_ATP_ trajectories of cell lineages under the indicated nutrient conditions. Cell divisions are represented by red vertical lines. (B-D) Distributions of single-cell growth rate, ATP level, and ATP fluctuation magnitude under different nutrient conditions. Data are represented in boxplots, with the medians labeled by the circles and 25%-75% percentiles shown by the square boxes.

### ATP fluctuations are attenuated in acetate fermentation mutants

We hypothesized that the differences in ATP fluctuations seen across growth media may originate, at least in part, from differences in the metabolic pathways used to generate ATP. In the presence of oxygen, cells are able to use aerobic respiration to generate ATP. However, under nutrient-rich conditions such as in M9GlcCAA, cells are also well-known to direct excess carbon molecules to less energy-efficient fermentation pathways, leading to secretion of fermented products outside the cell^32-35^. This is known as overflow metabolism.

In *E. coli*, there are three major fermentation pathways known as the lactate, ethanol, and acetate fermentation pathways^36^ (Figure 5A). Under aerobic conditions, acetate fermentation is the preferred fermentation pathway for the cell^37^. To test whether fermentation affects ATP dynamics, we performed microfluidic experiments in which we monitored the R_ATP_ of mutants defective in either acetate (Δ*pta* or Δ*ackA*), lactate (Δ*ldhA*) or ethanol (Δ*adhE*) fermentation while growing in M9GlcCAA. The lactate (Δ*ldhA*) and ethanol (Δ*adhE*) fermentation mutants had similar ATP dynamics and growth rate to the reference strain (Figure 5B and C), consistent with the idea that the lactate and ethanol fermentation pathways are largely inactive under aerobic condition. The average ATP level of the lactate mutant appeared slightly lower on average, though it remained within the variability range (Figure 5D). The ATP fluctuation level in these mutants was normal (Figure 5E).

**Figure 5:**
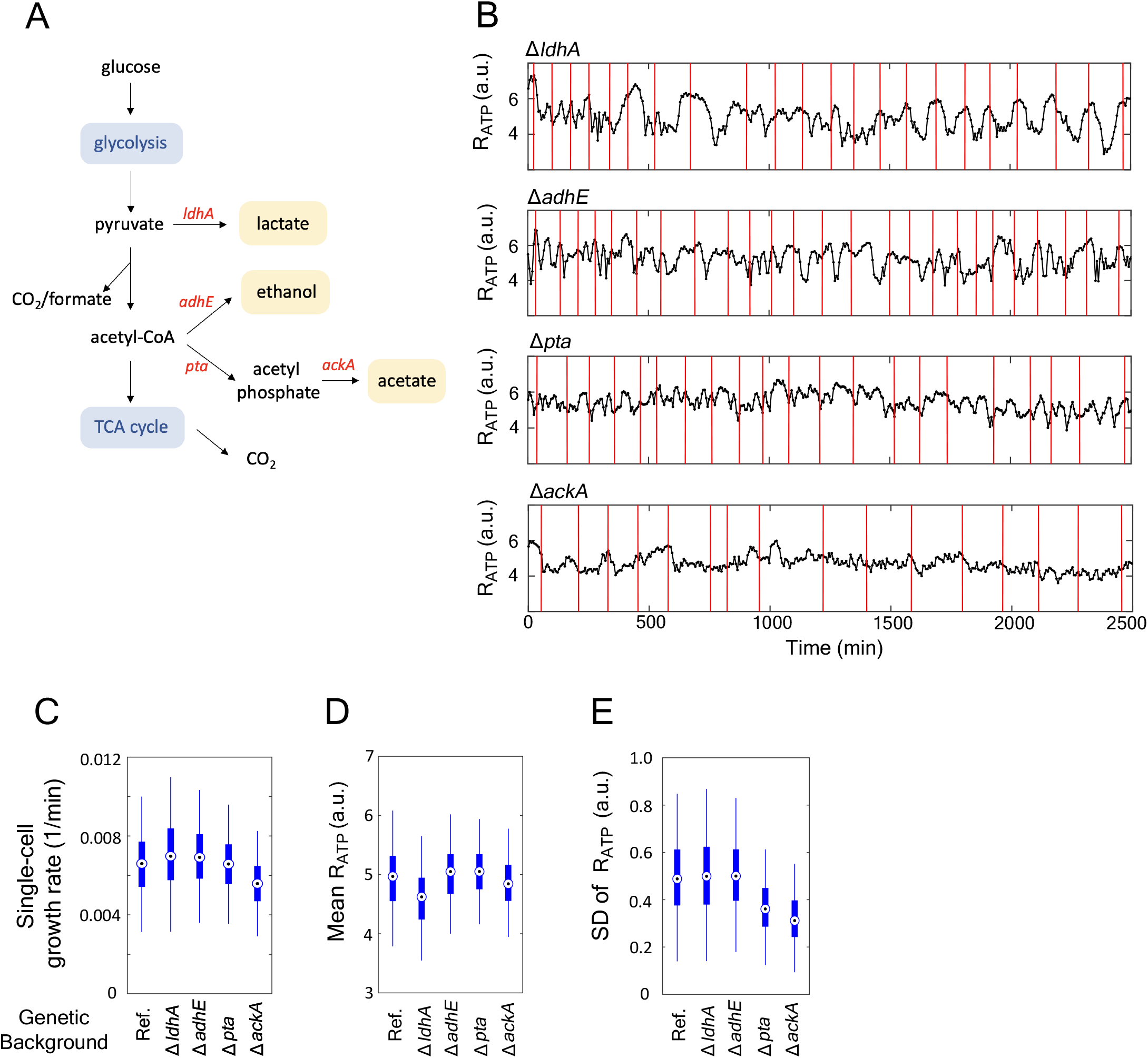
ATP dynamics of fermentation mutants growing in M9GlcCAA. (A) Schematic showing the *E. coli* fermentation pathways and the genes involved in them. TCA indicates tricarboxylic. (B-F) Data of the reference strain and the indicated fermentation mutants growing in M9GlcCAA. (B) Representative ATP trajectories of different fermentation mutants. Cell divisions are represented by red vertical lines. (C-E) Distributions of single-cell growth rate, ATP level, and ATP fluctuation magnitude for the reference strain and the indicated fermentation mutants. Data are shown as boxplots, with the medians labeled by circles and the 25%-75% percentiles shown by the boxes.

In contrast, the levels of ATP fluctuations were reduced by 30% to 40% in the acetate fermentation mutants (Δ*pta* or Δ*ackA*) while the mean ATP levels remained unaltered (Figure 5D and E). Notably, the reduced ATP fluctuation level in the acetate fermentation mutants was similar to that of wild-type cells growing under M9Xyl and M9Gly conditions (Figure 4B and D) under which acetate fermentation is expected to be considerably reduced or non-existent^38-40^. The lower ATP fluctuations in the acetate fermentation mutants was not simply due to a growth rate difference, as the Δ*pta* mutant had similar growth rate to the reference strain (Figure 5C). Our results suggest that in a rich nutrient condition, the process of acetate fermentation contributes to a significant portion (30-40%) of the ATP fluctuations.

### Acetate fermentation is associated with cell cycle-coordinated peaks of ATP levels near the division/birth time

In addition to increasing the level of ATP fluctuation, acetate fermentation also appears to contribute significantly to the preferred cell cycle phase of ATP fluctuations. We showed this by comparing the relative level of ATP (z-score of R_ATP_) along the cell cycle between the reference strain and the acetate fermentation mutants growing in M9GlcCAA. As discussed above, in the reference strain in which acetate fermentation occurs, the relative ATP level tended to peak around the times of division/birth (i.e., near 0% or 100% percent of the cell cycle) across many cell cycles (Figure 2D). This trend was reproduced in the ethanol and lactate fermentation mutants (Figure 6A). However, in the Δ*pta* and Δ*ackA* mutants, which are unable to carry out acetate fermentation, ATP peaks appeared to occur more random along the cell cycle (Figure 6A). To examine this quantitatively, for each cell cycle, we determined when the highest level of ATP occurs during the cell cycle and then averaged the results across cell cycles (Figure 6B). This analysis confirmed that that the ATP peak occurrence is higher around the time of cell division for the reference strain and the ethanol and lactate fermentation mutants. This enrichment was considerably reduced for the Δ*pta* and Δ*ackA* mutants. This suggests that ATP generated by acetate fermentation preferentially accumulates near division/birth. Consistent with this hypothesis, the enrichment in high z-score of R_ATP_ and ATP peak occurrence was reduced during growth on glucose and xylose, and disappeared during growth on glycerol (Figure 6A-B), reflecting the decreasing capability of acetate fermentation in these different media. These results suggest that acetate fermentation preferentially occurs near the beginning and end of the cell cycle in *E. coli*.

**Figure 6:**
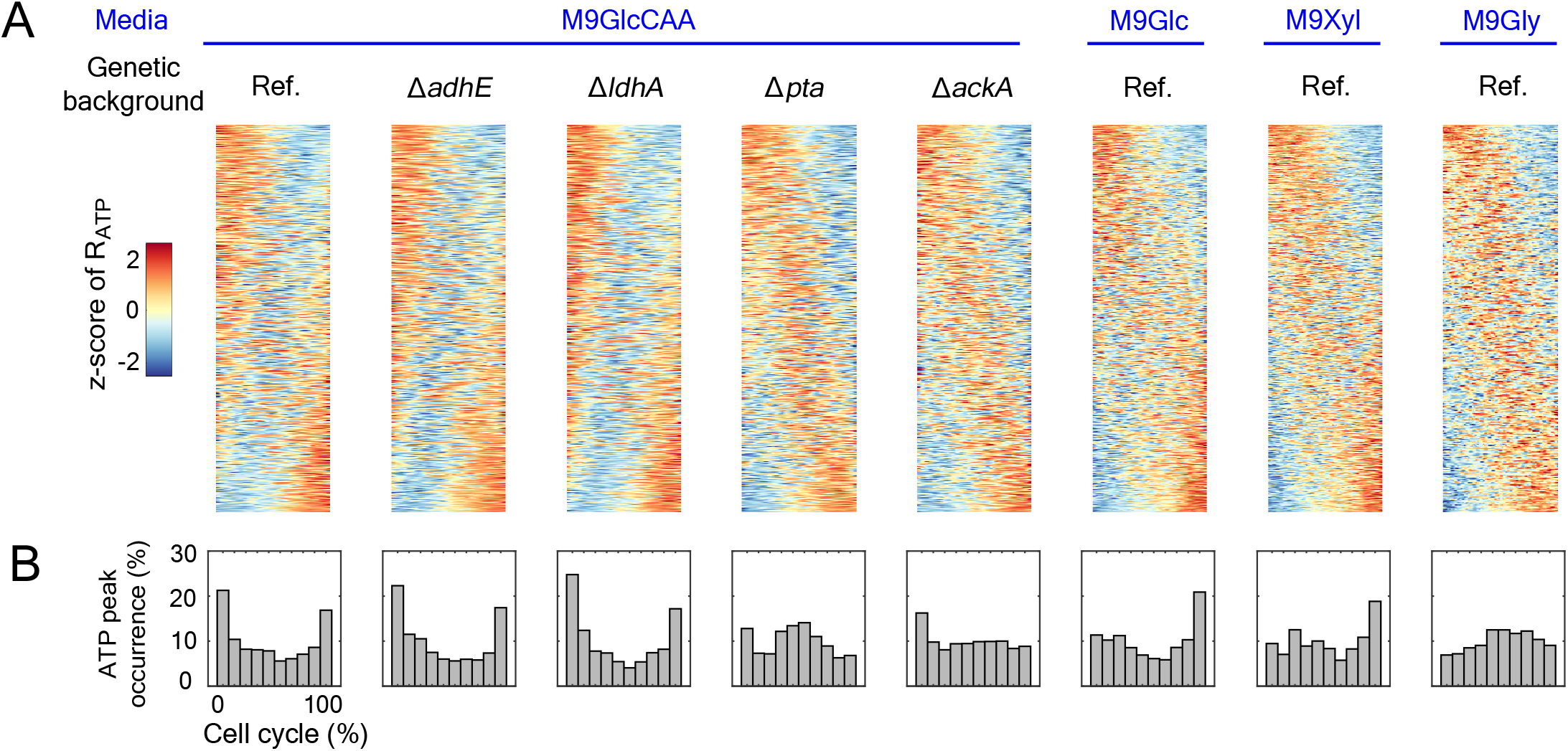
Cell-cycle tempograms of ATP patterning in different media and fermentation mutants. (A) Cell-cycle tempograms of z-transformed ATP pattern of different strains (or genotypes) in the indicated growth media. Each row in the tempogram represents one cell cycle (normalized to 0% - 100% of cell cycle percentile), with the z-transformed R_ATP_ level shown in the color scale. The rows in the tempogram are sorted according to the descending trend of R_ATP_ dynamics. The tempogram for M9GlcCAA is a replicate of Figure 2D. (B) Statistical distribution of R_ATP_ peak occurrence along the cell cycle (see STAR*Methods). For each cell cycle, the time bin with the highest R_ATP_ value was identified and referred to as the position of R_ATP_ peak.

### Smaller ATP fluctuations are associated with faster single-cell growth rates across tested metabolic conditions

We have shown that in M9GlcCAA, cells with smaller ATP fluctuations tend to have higher growth rates (Figure 2E, middle plot). We performed a similar analysis for all conditions tested (different media or genetic backgrounds). We found that the amplitude of ATP fluctuation negatively correlates with the single-cell growth rate in all conditions (Figure 7A), though this negative correlation was small for the glycerol case for which the growth rate and ATP fluctuations remain low across cells. Collectively, this negative correlation suggests that tight ATP homeostasis may be beneficial across metabolic conditions. Whether the cells grow in nutrient-rich or nutrient-poor media, performing overflow metabolism or not, they tend to grow better when their ATP levels are maintained close to a constant.

**Figure 7:**
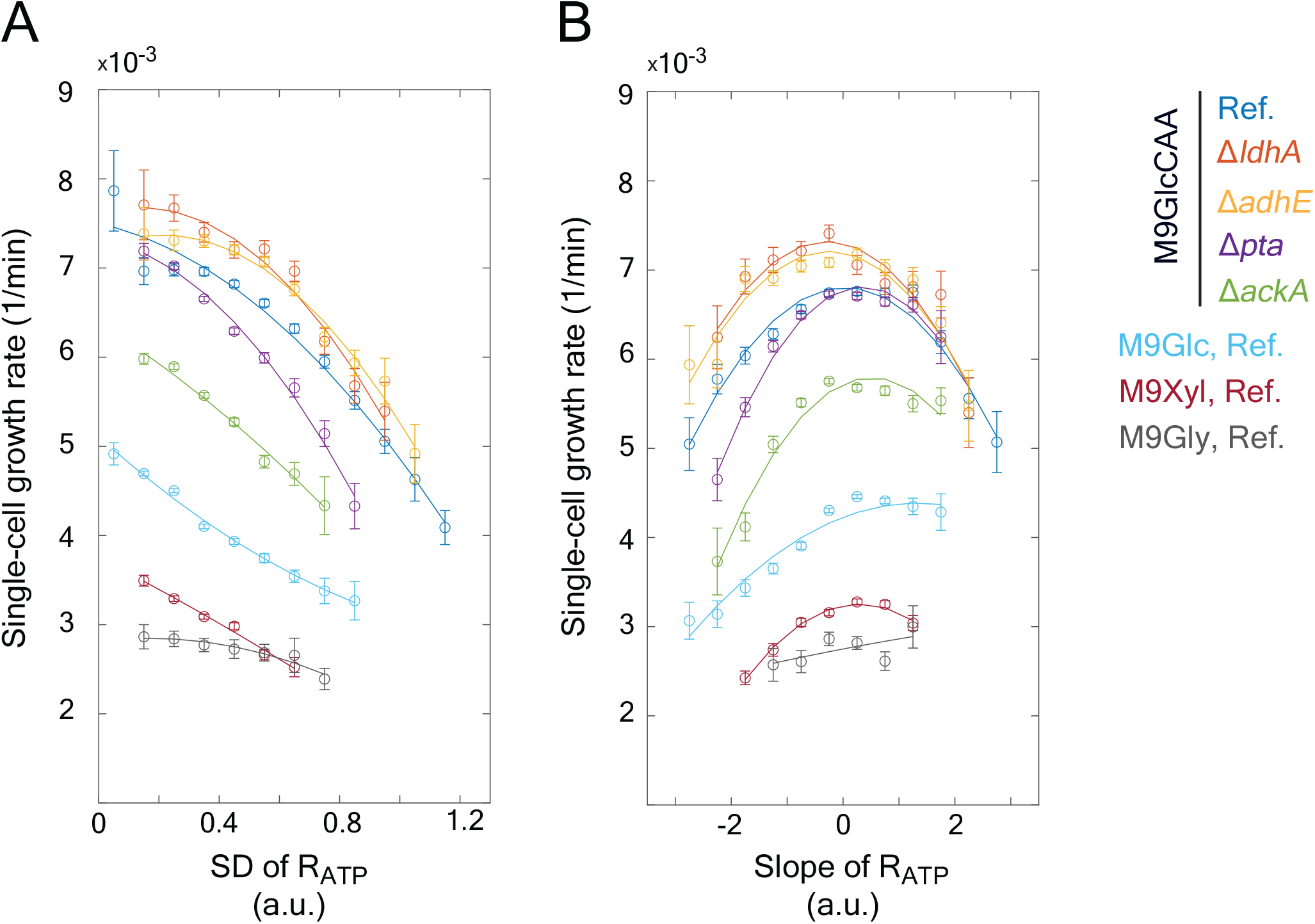
Correlations between ATP fluctuation pattern and single-cell growth rate. (A) Plot showing the single-cell growth rate as a function of the SD of R_ATP_. (B) Plot showing the single-cell growth rate as a function of the slope of the R_ATP_ trajectory (see schematic in Figure 2C). For both panels A and B, each strain and media condition are depicted in a different color according to the legend at the right.

Similarly, the correlation between growth rate and the slope of the ATP level during the cell cycle (Figure 7B) tended to follow an inverted U-shape across virtually all conditions (except again for the glycerol condition for which there was too little variability for either variable to be informative). These results indicate that cells have a similar growth rate reduction whether they gradually increase or decrease their ATP concentration during the cell cycle.

## Discussion

Our single-cell study in *E. coli* reveals several important aspects of ATP dynamics and homeostasis in *E. coli* with respect to overflow metabolism, the cell cycle, and cell growth.

### Overflow metabolism and ATP fluctuation

Overflow metabolism, also referred to as the Warburg or Crabtree effect, is widely observed in fast-growing bacteria and yeast, as well as in cancer cells^32-34,41-45^. It is thought to be a metabolic strategy by which cells maximize biomass synthesis under nutrient-rich conditions by using proteomic-efficient aerobic fermentation over carbon-efficient respiration^35^. To our knowledge, our study provides the first insight into overflow metabolism at the single-cell level. We find that under condition of overflow metabolism (M9GlcCAA), the ATP level in *E. coli* fluctuates considerably around the estimated mean value of 2.4 mM during the cell cycle, with an average peak and valley of 3.6 mM and 1.2 mM, respectively. Shutting down overflow metabolism by deletion of genes involved in acetate fermentation results in reduced ATP fluctuations by about 25% - 35% (Figure 5B and E). Growth in the presence of carbon sources that reduce overflow metabolism is also accompanied by reduced ATP fluctuations by 27% - 36% (Figure 4A and D). These observations argue that production of ATP via aerobic acetate fermentation causes large fluctuations in ATP concentrations inside *E. coli* cells.

How could this happen? While answering this question will require further study, the answer may be linked to two observations: (i) peaks in ATP concentration are enriched around the end of a cell cycle and the beginning of the next one, under conditions favoring aerobic acetate fermentation (Figure 2D), and (ii) this enrichment in ATP peaks around the time of division decreases or disappears under culture and genetic conditions that diminish or abolish acetate fermentation (Figure 6). This suggests that aerobic acetate fermentation (or overflow metabolism) does not occur homogeneously throughout the cell cycle and is instead more active around the time of cell division (Figure 6). A preferential boost in ATP synthesis during a specific time window will result in ATP accumulation if the ATP consumption (i.e., the metabolic demand) is not matched.

### Temporal ATP dynamics and the cell cycle

What regulates the temporal dynamics of ATP (or acetate fermentation/overflow metabolism) in *E. coli* is not intuitive. Interestingly, energy metabolism in eukaryotic cells is linked to the cell cycle via a variety of signaling pathways that involve different regulators or processes such as the AMP-activated protein kinase, mTOR, cyclins, cyclin-dependent kinase inhibitors, histone acetylation, and ubiquitin-dependent degradation ^14-18,46 19-21,47-49^. None of these proteins or processes are present in bacteria. Yet, we identify an ATP oscillatory pattern with a cell cycle-like period in *E. coli* (Figure 2A-D), which occurs about 40-50% of the time based on our wavelet analysis of cell lineages. This may highlight a basic cellular principle that promotes (at least partial) synchronization between energy metabolism and cell cycle progression. During evolution, addition of sophisticated regulatory mechanisms, as those observed in today’s eukaryotic organisms, may have increased the robustness in the coupling between energy production and the cell cycle.

### ATP fluctuations and cell growth

It is well-known that under a given growth condition, isogenic cells display considerable variability in growth rate^23,24^. The molecular origin of this variability is poorly understood. Our study identifies ATP fluctuations as a potential contributor to cell growth heterogeneity in a clonal population. This is suggested by the striking negative correlation between ATP fluctuation and growth rate at the single-cell level (Figure 7A). That is, for a given growth condition, cells that display a large change in ATP concentration during the cell cycle tend to grow slower than cells with smaller ATP fluctuations. This negative correlation is robust to differences in strains or carbon sources used in the study (Figure 7A), suggesting the existence of a meaningful relation between ATP fluctuations and cell growth rate. Dilution by growth (i.e., expansion of the cellular volume) cannot explain this relation as the negative correlation persists regardless of whether the concentration in ATP is descending or ascending during the cell cycle (Figures 2E and 7B). Instead, the fastest (optimal) growth is associated with the most stable (constant) ATP concentration.

It is possible that the negative correlation is not causal. Perhaps stress or other unknown factors affect the stability of the ATP level and the growth rate of the cell independently. Even if that were true, understanding the origin of this indirect connection is likely to help us understand how cells control their growth and ATP production or consumption. A more attractive hypothesis is that large ATP fluctuations are detrimental to cell growth. For example, cells may have to dynamically adjust their metabolism in response to fluctuating ATP levels. There might be a cost associated with bringing the system back to the optimal level, potentially through regulation of energy storage (e.g., in the form of polyphosphate and glycogen). Broadly speaking, this would be analogous to electric grid systems in which fluctuating power demand and intermittent energy production (e.g., from solar and wind output) are associated with costs, notably for energy storage management^50^. Another mutually non-exclusive possibility may be linked to the recently proposed function of ATP as a biological hydrotrope that help solubilize proteins in the cytoplasm^51^. Perhaps sudden changes in cellular ATP concentration have detrimental effects on protein solubility, which may lead to a growth penalty. Other possibilities likely exist.

While these ideas remain speculative, our study highlights the importance of time-resolved single-cell studies, even for molecules as abundant and stable in concentration at the population level as ATP. ATP may not be the only metabolite whose levels fluctuate in bacteria. A recent single-cell study has proposed that the redox cofactors NADH and NADPH oscillate during the cell cycle in *E. coli* based on cellular autofluorescence measurements ^52^. Future quantitative single-cell studies will be facilitated with the recent exciting development of various biosensors that sense the concentration of different metabolites at physiological levels^53^. This may have broad implications not only from a basic science standpoint but from a metabolic engineering perspective, as current metabolic models generally assume that cells are under balanced growth, where metabolites (including ATP) are maintained at a constant level. Expanding these models beyond balanced growth, for instance, by implementing a scalable reaction network approach^54,55^, may offer improved predictive strategies for strain design.

## STAR*Methods

### Key resource table

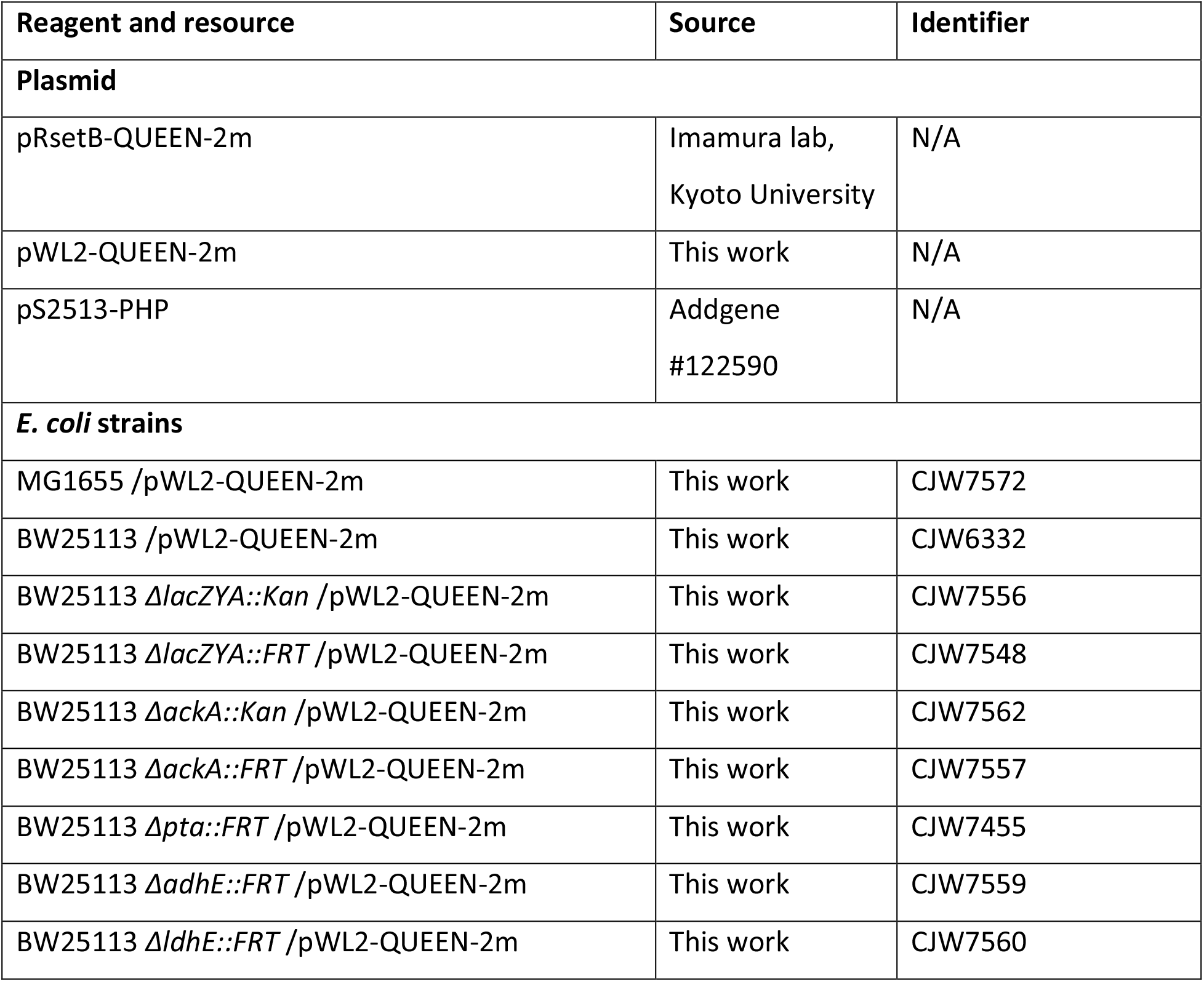
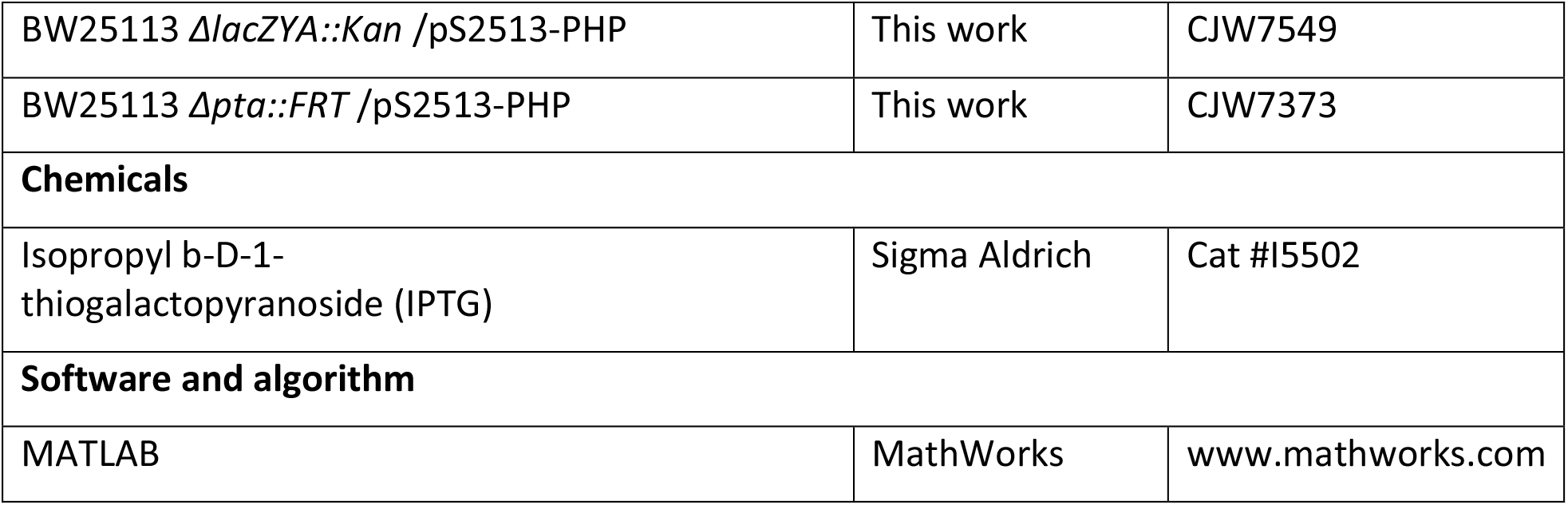

### Resource availability

#### Lead contact

Requested for resource and reagents and further information should be directed to and be fulfilled by the lead contact, Christine Jacobs-Wagner.

#### Materials availability

Strains and plasmids generated in this study are available without restriction from the lead contact Christine Jacobs-Wagner.

#### Data and code availability

The MATLAB script for data analysis and the processed dataset can be found on the following Github website: https://github.com/JacobsWagnerLab/.

### Experimental model and subject details

#### Strains, constructs and growth conditions

To achieve tunable expression of the QUEEN-2m ATP biosensor, plasmid pWL2 was constructed for the general purpose of gene expression. pWL2 was obtained by introducing a DNA fragment from pMMB67EH (including the *tac* promoter, a multiple cloning site, and the *rrnB* terminator) into pACYCDuet1, which contains a *lacI* repressor gene. The *tac* promoter, a fusion of *trp* and *lac* promoters, can be derepressed by addition of Isopropyl β-D-1-thiogalactopyranoside (IPTG) while being less sensitive to glucose inhibition than the *lac* promoter^56^. The open reading frame of the QUEEN-2m biosensor (acquired from the Imamura lab, Kyoto University^22^) was cloned into the multiple cloning site of pWL2, thus generating plasmid pWL2-QUEEN-2m. This construct allowed us to express the QUEEN-2m biosensor during growth in various glucose-containing media.

Our study used two reference strains, CJW5187 and CJW7453. CJW5187 was constructed by replacing the *lacZYA* operon in the Keio parental strain (BW25113) with a kanamycin resistance cassette (KanR). This strain serves as the reference for the Keio collection mutants^57^, all of which have a specific gene replaced by KanR. The second reference strain, CJW7453, was generated by flipping out the KanR in CJW5187 by using the FLP recombination method ^57^. The two reference strains (with and without KanR) display similar ATP dynamics (Figure S4). Therefore, the two datasets were combined for presentation (Figures 2,3,4,5,6, and 7) and further analysis. To determine the protentional role of fermentation pathways in ATP dynamics, we acquired the Δ*ackA*, Δ*pta*, Δ*adhE*, and Δ*ldhA* strains from the Keio collection, in which *ackA, pta, adhE*, or *ldhA* are substituted by KanR. Corresponding mutant strains without KanR (strains CJW7454, CJW7455, CJW7546, CJW7547 for Δ*ackA*, Δ*pta*, Δ*adhE*, and Δ*ldhA*, respectively) were generated by flipping out KanR. The ATP dynamics of the Δ*ackA* strains, either with or without the KanR, were similar (Figure S4). Therefore, the two datasets were combined for presentation (Figures 5,6, and 7) and further analysis.

The plasmid pWL2-QUEEN-2m was introduced into the reference and fermentation mutant strains by heat shock transformation, creating CJW7556 and CJW7548 for the reference strains with and without KanR, CJW7562 and CJW7557 for Δ*ackA* with and without KanR, and CJW7558, CJW7559 and CJW7560 for Δ*pta*, Δ*adhE*, and Δ*ldhA* without KanR, respectively. Unless indicated, 60 μM of IPTG was used to induce the synthesis of the QUEEN-2m biosensor. To measure pH dynamics, plasmid pS2513-PHP^28^ (Addgene #122590) was introduced into the reference strain CJW5187 by heat shock transformation, yielding strain CJW7549 in which the pH sensor protein (PHP) is constitutively expressed by the EM7 promoter^28^.

### Method details

#### Fabrication of the microfluidic devices

The master mold of our microfluidic devices was manufactured in the clean room of the School of Engineering and Applied Science at Yale University. UV photolithography (EVG 620 Mask Aligner) was applied on Si wafers with SU-8 photoresist to generate two layers of positive patterns. The lower layer contained the bacterial culture chambers while the upper layer contained the large trench for flowing culture medium. A detailed protocol on the photolithography and design of the microfluidic device is provided in Supplementary Information. The polydimethylsiloxane (PDMS) (Sylgard 184, Dow) casts, which has negative microfluidic patterns, was generated from the Si wafer with positive SU-8 patterns. After punching the inlets and outlets using a biopsy needle of 1.2 μm diameter, the PDMS replica and glass-bottom Petri dish were treated with air plasma (Harrick Plasma PDC-32G) and bonded together, creating the microfluidic device. To strengthen the bonding, the microfluidic devices were placed at 65°C for 20 min.

#### Microfluidic experiments

The M9 buffer used in this study contains 3.4 mM of Na_2_HPO_4_, 2.2 mM of KH_2_PO_4_, 0.85 mM of NaCl, 0.93 mM of NH_4_Cl, 1.0 mM of Mg_2_SO_4_, 0.3 mM of CaCl_2_, 0.1%(w/w) of thiamine, and trace elements containing 134 μM of ethylenediaminetetraacetic acid (EDTA), 31 μM of FeCl_3_-6H_2_O, 6.0 μM of ZnCl_2_, 0.8 μM of CuCl_2_-2H_2_O, 0.4 μM of CoCl_2_-2H_2_O, 1.6 μM of HBO_3_-2H_2_O, 0.8 μM of MnCl_2_-4H_2_O. Culture media used in the experiments were M9GlcCAA (M9 buffer with 0.2% (w/v) glucose and 0.5% (w/v) of casamino acid), M9Glc (M9 buffer with 0.2% (w/v) glucose), M9Xyl (M9 buffer with 0.2% (w/v) xylose), and M9Gly (M9 buffer with 0.2% (v/v) glycerol). Cells were grown in aerated liquid cultures that maintained exponential growth for at least 12 h at 25°C in 200 rpm shaking water bath under constant IPTG induction of QUEEN-2m expression. Then, cells were concentrated 100 times by centrifugation (6000 rpm for 10 min) and resuspended in the appropriate growth medium prior to injection into the microfluidic device. For the stationary phase experiments, cells were cultured in M9GlcCAA in the presence of IPTG until the culture reached stationary phase. The cells were then concentrated by centrifugation and resuspended in M9 buffer where they remained under shaking conditions for 2 h before cell injection into the microfluidic device.

Before cell injection, the microfluidic device was blocked with 5% BSA dissolved in the appropriate culture medium. Resuspended cells were blocked with 10% BSA for 2 min prior to injection into the microfluidic device. The microfluidic device was then mounted on the microscope for visual inspection. After cells entered the chambers, fresh medium was connected to the microfluidic device. Cells were then allowed to recover and grow for 3 h prior to image acquisition.

For medium switching experiments, a customized setup similar to one previously described^30^ was used. Briefly, μ-Manager scripts (https://github.com/JacobsWagnerLab/) were used to program the Arduino Uno microcontroller (Arduino), which triggered the opening and closing of solenoid valves (Cole-Parmer) and thereby controlled the flow of the culture media. The media flows were driven by constant pressure (10-20 psi) such that the turnover rate of growth media in the microfluidic device was about 5 times/s.

### Microscopy

Time-lapse imaging was done using a Nikon Eclipse Ti microscope equipped with a Hamamatsu ORCA-Flash 4.0 camera, a 100X objective (Nikon, OFN25 Ph3 DM, N.A. 1.45) and type N lens oil (Nikon). Unless indicated, the temperature was maintained at 25 ± 0.2°C using a customized enclosure and a temperature controller (Air-Therm SWT, World Precision Instrument). Phase contrast and fluorescence images were acquired every 1 and 6 min, respectively. Under our imaging condition, 1 μm^2^ area corresponds to about 272 pixels.

Ratiometric imaging of QUEEN-2m and PHP biosensors was done using the violet (wavelength = 395 ± 25 nm) and cyan (wavelength = 470 ± 24 nm) LED light sources (Spectra X, Lumencor) for the two excitation wavelengths of the biosensors. Fluorescence imaging was performed using a triple band filter set (405/488/561nm) (#69901, Chroma).

### Conversion of R_ATP_ values into absolute ATP concentrations

See Supplementary Information for a detailed description of the microfluidic fabrication, the luciferase assay and the method used to generate a calibration curve that converts R_ATP_ values into intracellular ATP concentrations.

### Image and data analysis

A customized MATLAB-based image analysis pipeline was used to analyze the microfluidic data. The pipeline consists of multiple parts, including (1) shift correction for image stacks, (2) cropping region of interest (ROI) and removal of microfluidic background features (3) cell segmentation based on adaptive threshold algorithms, (4) cell segmentation refinement and cell division detection based on a kymograph algorithm, (5) cell lineage tracking, (6) background subtraction from the fluorescence images, and (7) cell area and fluorescence intensity measurements of individual cells. The MATLAB code for the pipeline can be found at https://github.com/JacobsWagnerLab.

To calculate the R_ATP_, the fluorescence values of 405 nm or 488 nm excitation channels were averaged for all pixels in each cell. The background fluorescence in the microfluidic device was subtracted from the average values for both channels. The R_ATP_ was defined by the ratio between the mean fluorescence intensities at 405 nm and 488 nm. Since the R_ATP_ is calculated for individual cells, the mean 405nm and 488nm intensity values were averaged from more than 200 pixels, resulting in a negligible measurement error from the CMOS camera noise. The R_ATP_ is also robust to small differences in cell contours, since the R_ATP_ measurements are ratiometric and both fluorescence intensity measurements are similarly affected by a small change in the cell boundary region. The level of QUEEN-2m protein was inferred from analyzing the signals from the 405 nm and 488 nm excitation channels; see Supplementary Information for a detailed description.

The calculation of R_pH_ was done in the same way as for the R_ATP_. To estimate the absolute fluctuation of pH, the coefficient of variation (CV) of our R_pH_ measurements was calculated to be about 2.5%. From the original description of the PHP sensor^28^, a 2.5% CV of R_pH_ at pH =7.5 (the typical pH in the *E. coli* cytosol) corresponds roughly to a pH change of ±0.05 pH units.

To calculate the cell cycle growth rate, the log-transformed cell area data were fitted with a linear function and the slope was used to define the cell cycle growth rate. Cell-cycle tempograms were constructed by collecting R_ATP_ (or R_pH_) data of all cell cycles. For each cell cycle, the data were interpolated from 0% to 100% of cell cycle percentile and presented in a row vector. For each cell cycle, the time point with the highest R_ATP_ value was identified and referred to as the position of R_ATP_ peak. The R_ATP_ peak occurrence was calculated for each time bin (10% of the cell cycle).

For the cell lineage analysis, all values of R_ATP_ (or R_pH_) were collected along the lineage of a mother cell (the cell located at the terminal position of the microfluidic chambers) and catenated into a data vector. For wavelet analysis, the baseline of the data vector was first determined using the Elementary Mode Decomposition (EMD) method^58^, which decomposes the data vectors into oscillatory modes of different time scales. The modes with mean wavelengths at least twice longer than the mean cell cycle time were regarded as components of the baseline. Once the baseline signal was calculated, the detrended data vector (with the baseline removed) was used for wavelet analysis. The MATLAB built-in functions *emd* and *cwt* (with ‘amor’ (Gabor) wavelet type) were used for EMD analysis and wavelet analysis, respectively.

## Acknowledgements

We are grateful to Dr. Hiromi Imamura for the gift of the QUEEN ATP sensor constructs. We thank Dr. Michael Power, director of Yale University Cleanroom for helping us to design and manufacture the microfluidic devices used in this study. We thank Dr. Yi-Hao Kao for valuable comments on time-series analysis. We thank all the other members of the Jacobs-Wagner laboratory for fruitful discussion and critical reading of the manuscript. C.J.-W.is an investigator of the Howard Hughes Medical Institute.

## Author Contributions

W.-H.L. and C.J.-W. designed the study. W.-H.L. performed the experiments, established the data analysis pipeline, and analyzed the data. W.-H.L. and C.J.-W. wrote the manuscript. C. J.-W. supervised the study and secured the funding.

## Declaration of interests

The authors declare no competing interests.

**Figure S1: R**_**ATP**_ **dynamics and corresponding biosensor protein levels during cell growth in different media**

Representative data of R_ATP_ (upper panel) and QUEEN-2m protein level (lower panel) in cell lineages in the indicated growth medium. Red vertical lines represent cell divisions.

**Figure S2: ATP dynamics measured at different levels of the biosensor protein QUEEN-2m expression**

In this study, the typical concentration of IPTG used to induce the expression of the biosensor QUEEN-2m was 60 μM; see Figure 2. Here, the effect of reducing the level of biosensor in the cell by inducing its expression with only 15 μM was examined. The growth medium was M9GlcCAA. (A) Distributions showing the protein and protein fluctuation level of the biosensor under two different levels of IPTG induction. The data are represented in boxplots, with the medians labeled by the circles and the 25%-75% percentiles shown by the square boxes. (B) Distributions of growth rate, ATP level, and ATP fluctuation level at different concentrations of IPTG induction. (C) Representative R_ATP_ dynamics in a cell lineage when 15 μM IPTG was used as induction condition. (D) Wavelet analysis scalogram of (C). (E) cell cycle pattern of R_ATP_ (absolute values) for all collected cell cycles under different biosensor protein levels. (F) Similar to (E), except with z-transformed R_ATP_ for every cell cycle. (G) Comparison of correlation between single-cell growth rate and different ATP statistics (mean R_ATP_ level, SD of R_ATP_, or slope of R_ATP_ during the cell cycle) under different conditions of biosensor protein expression (15 vs. 60 μM IPTG induction).

**Figure S3: pH dynamics during exponential growth**

The data were collected using the fluorescence pH sensor PHP expressed in *E. coli* cells (strain CJW7549) growing in M9GlcCAA medium. (A) pH fluctuation during normal exponential growth. Upper panel: Trajectory of R_pH_ using the pHluorin2 pH sensor. Lower panel: Wavelet analysis scalogram of the upper panel. (B) Similar to (A), except using the acetate fermentation mutant *Δpta*. (C) Distribution of single-cell growth rate, R_pH_ (proxy for pH level), and R_pH_ fluctuation level (proxy for pH fluctuation level) of the reference strain and fermentation mutant *Δpta*. (D) Cell cycle tempogram of pH dynamics. Each row represents one cell cycle, with pH level (using R_pH_ as a proxy) labeled in color (see Figure 2C).

**Figure S4: ATP dynamics in different *E. coli* strain background**

The data were collected with different genetic backgrounds of *E. coli* expressing plasmid encoded QUEEN-2m and growing in M9GlcCAA, with n = 1700 to 3000 cell cycles for each strain. (A) Distributions of single-cell growth rate, R_ATP_, and SD of R_ATP_ for the indicated strains. Data are shown in boxplots, with the medians labeled by the circles and 25%-75% percentiles shown by the square boxes. (B) Cell-cycle tempograms of ATP of each indicated strain. Each row in the tempogram represents one cell cycle (normalized to 0% - 100% of cell cycle percentile), with the z-transformed R_ATP_ level shown in the color scale. The rows in the tempogram are sorted according to the descending trend of R_ATP_ dynamics. (C) Statistical distribution of R_ATP_ peak occurrence along the cell cycle.

**Figure S5: Growth rate and ATP statistics for the cell cycles of different cell ages and positions in the microfluidic device**

The data were collected in M9GlcCAA medium. (A) Schematics showing the pole classes from 1 to 4 in a cell lineage, with the ages of cell poles being labeled 0, 1, 2,…, n, n+1, n+2 from youngest to oldest. The cell cycles of the cells located at the terminal position of the microfluidic chamber belong to class 1. (B) Statistics for different cell cycles belonging to the pole classes defined in (A).

**Figure S6: Distributions of coefficients of variation (CV) for ATP concentration in different media and fermentation mutants**.

(A) The CV of the calibrated ATP concentration for the reference strain in different media. (B) The CV of the calibrated ATP concentration for the indicated genetic backgrounds in M9GlcCAA medium. Data are represented in boxplots, with the medians labeled by the circles and 25%-75% percentiles shown by the square boxes.

**Movie S1: ATP level dynamics in *E. coli* cells**.

The movie shows two representative microfluidic chambers with *E. coli* cells (CJW7548; the reference strain) expressing the QUEEN-2m ATP sensor. Cells were grown in the presence of M9GlcCAA at 25°C. The ratiometric measurement R_ATP_ is shown with the color bar, where red to blue colors represent high to low values of R_ATP_. The time stamp indicates hh:mm, and the scale bar indicates 5 μm.

