## Supplementary Information for "Connecting single-cell ATP dynamics to overflow metabolism, cell growth and the cell cycle in *Escherichia coli*"

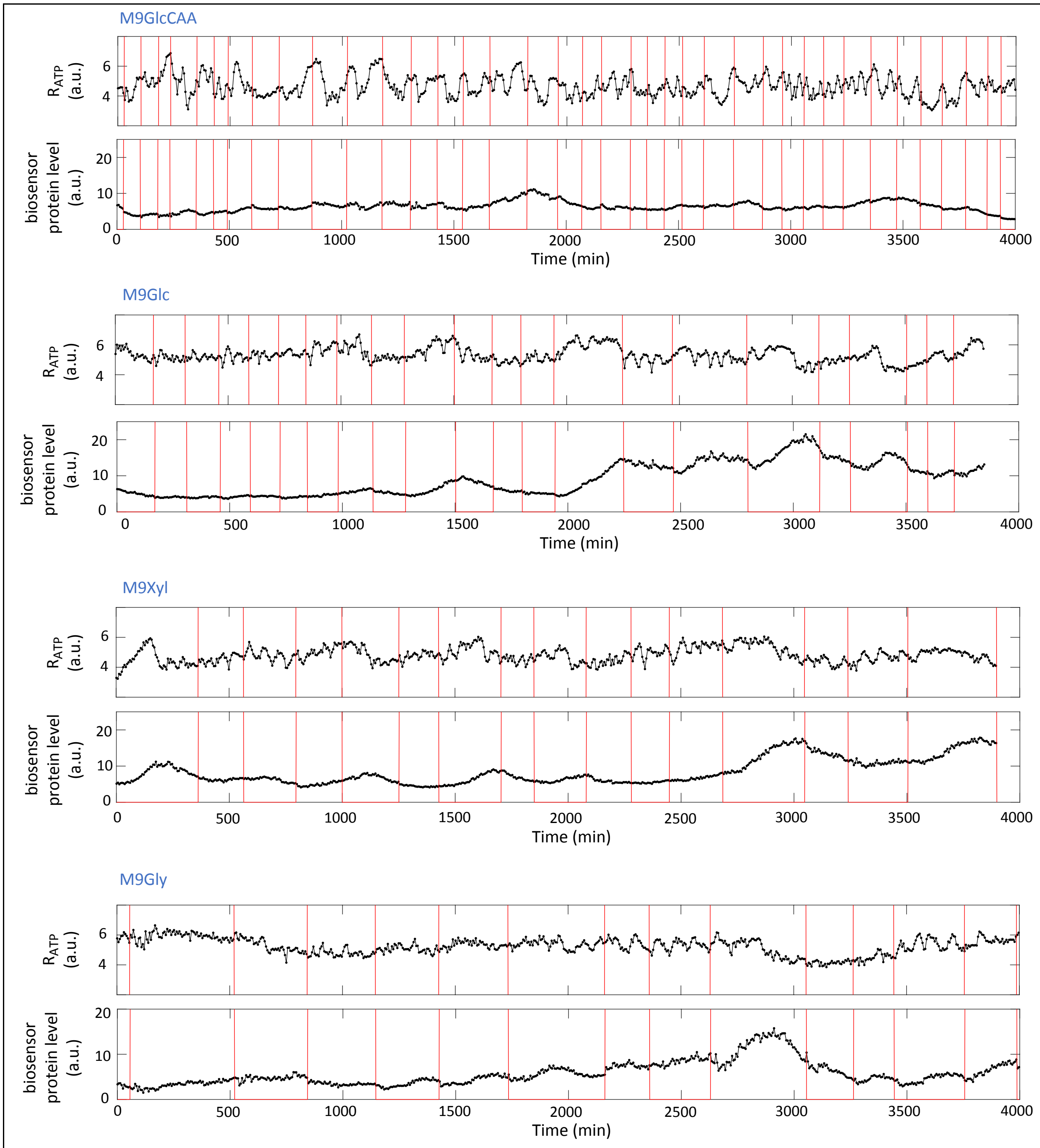


**Figure S1: $R_{ATP}$ dynamics and corresponding biosensor protein levels during cell growth in different media. Related to Figure 1.**

Representative data of $R_{ATP}$ (upper panel) and QUEEN-2m protein level (lower panel) in cell lineages in the indicated growth medium. Red vertical lines represent cell divisions.

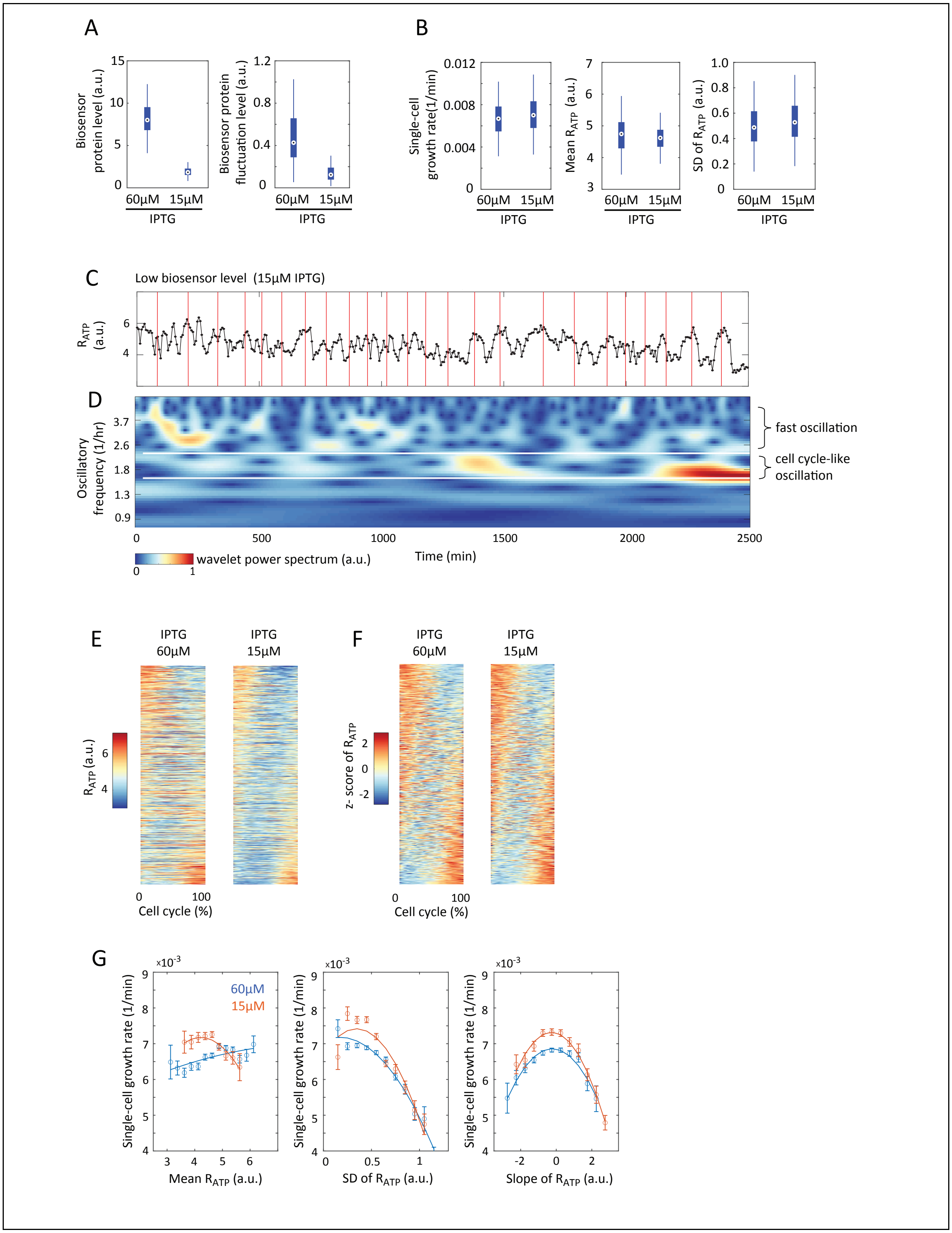
A
Biosensor protein level (a.u.)
Biosensor protein fluctuation level (a.u.)
60μM 15μM
IPTG
B
Single-cell growth rate(1/min)
Mean $R_{ATP}$ (a.u.)
SD of $R_{ATP}$ (a.u.)
60μM 15μM
IPTG
C
Low biosensor level (15μM IPTG)
$R_{ATP}$ (a.u.)
D
Oscillatory frequency (1/hr)
fast oscillation
cell cycle-like oscillation
wavelet power spectrum (a.u.)
Time (min)
E
IPTG 60μM
IPTG 15μM
$R_{ATP}$ (a.u.)
Cell cycle (%)
F
IPTG 60μM
IPTG 15μM
z- score of $R_{ATP}$
Cell cycle (%)
G
60μM
15μM
Single-cell growth rate (1/min)
Mean $R_{ATP}$ (a.u.)
SD of $R_{ATP}$ (a.u.)
Slope of $R_{ATP}$ (a.u.)

**Figure S2: ATP dynamics measured at different levels of the biosensor protein QUEEN-2m expression. Related to Figure 1.**

In this study, the typical concentration of IPTG used to induce the expression of the biosensor QUEEN-2m was 60 µM; see Figure 2. Here, the effect of reducing the level of biosensor in the cell by inducing its expression with only 15 µM was examined. The growth medium was M9GlcCAA. (A) Distributions showing the protein and protein fluctuation level of the biosensor under two different levels of IPTG induction. The data are represented in boxplots, with the medians labeled by the circles and the 25%-75% percentiles shown by the square boxes. (B) Distributions of growth rate, ATP level, and ATP fluctuation level at different concentrations of IPTG induction. (C) Representative $R_{ATP}$ dynamics in a cell lineage when 15 µM IPTG was used as induction condition. (D) Wavelet analysis scalogram of (C). (E) cell cycle pattern of $R_{ATP}$ (absolute values) for all collected cell cycles under different biosensor protein levels. (F) Similar to (E), except with z-transformed $R_{ATP}$ for every cell cycle. (G) Comparison of correlation between single-cell growth rate and different ATP statistics (mean $R_{ATP}$ level, SD of $R_{ATP}$, or slope of $R_{ATP}$ during the cell cycle) under different conditions of biosensor protein expression (15 vs. 60 µM IPTG induction).

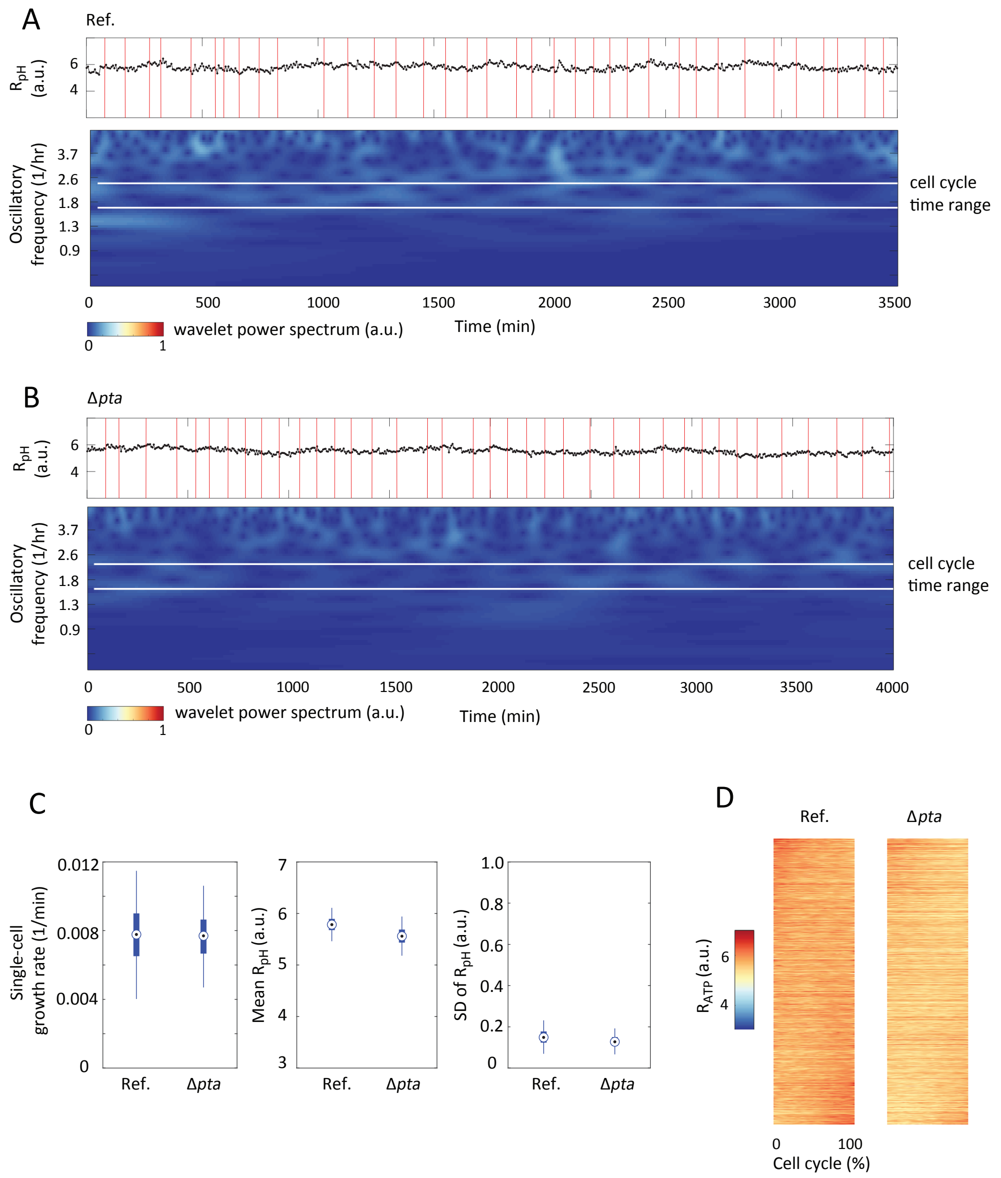


**Figure S3: pH dynamics during exponential growth. Related to Figure 1.**

The data were collected using the fluorescence pH sensor PHP expressed in *E. coli* cells (strain CJW7549) growing in M9GlcCAA medium. (A) pH fluctuation during normal exponential growth. Upper panel: Trajectory of $R_{pH}$ using the pHluorin2 pH sensor. Lower panel: Wavelet analysis scalogram of the upper panel. (B) Similar to (A), except using the acetate fermentation mutant *Δpta*. (C) Distribution of single-cell growth rate, $R_{pH}$ (proxy for pH level), and $R_{pH}$ fluctuation level (proxy for pH fluctuation level) of the reference strain and fermentation mutant *Δpta*. (D) Cell cycle tempogram of pH dynamics. Each row represents one cell cycle, with pH level (using $R_{pH}$ as a proxy) labeled in color (see Figure 2C).

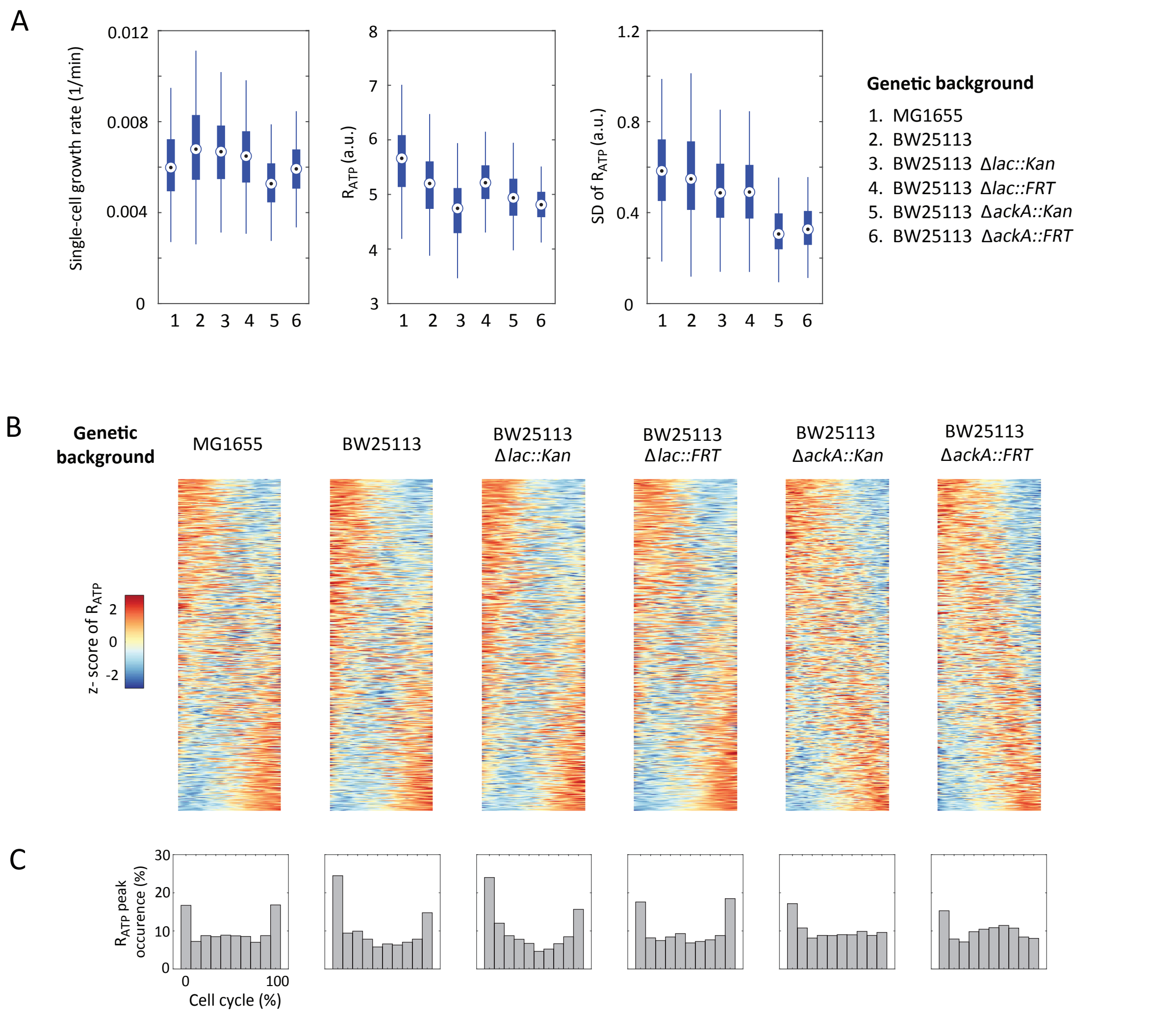


**Figure S4: ATP dynamics in different *E. coli* strain background. Related to Figure 1.**

The data were collected with different genetic backgrounds of *E. coli* expressing plasmid encoded QUEEN-2m and growing in M9GlcCAA, with n = 1700 to 3000 cell cycles for each strain. (A) Distributions of single-cell growth rate, $R_{ATP}$, and SD of $R_{ATP}$ for the indicated strains. Data are shown in boxplots, with the medians labeled by the circles and 25%-75% percentiles shown by the square boxes. (B) Cell-cycle tempograms of ATP of each indicated strain. Each row in the tempogram represents one cell cycle (normalized to 0% - 100% of cell cycle percentile), with the z-transformed $R_{ATP}$ level shown in the color scale. The rows in the tempogram are sorted according to the descending trend of $R_{ATP}$ dynamics. (C) Statistical distribution of $R_{ATP}$ peak occurrence along the cell cycle.

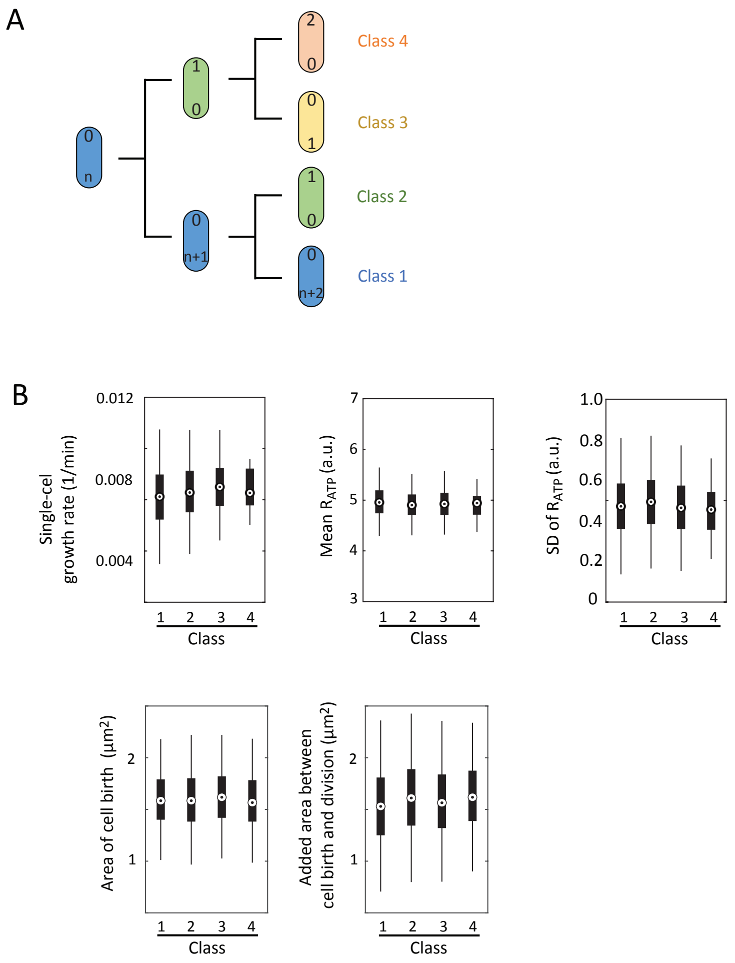


**Figure S5: Growth rate and ATP statistics for the cell cycles of different cell ages and positions in the microfluidic device. Related to Figure 3.**

The data were collected in M9GlcCAA medium. (A) Schematics showing the pole classes from 1 to 4 in a cell lineage, with the ages of cell poles being labeled 0, 1, 2,…, n, n+1, n+2 from youngest to oldest. The cell cycles of the cells located at the terminal position of the microfluidic chamber belong to class 1. (B) Statistics for different cell cycles belonging to the pole classes defined in (A).

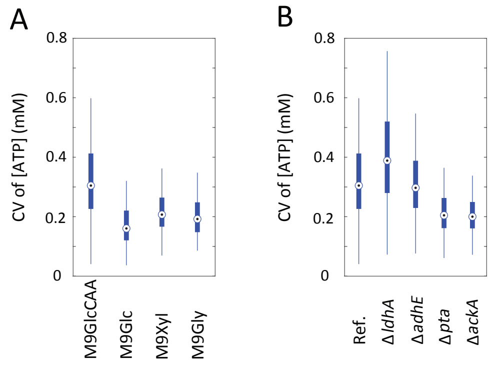


**Figure S6: Distributions of coefficients of variation (CV) for ATP concentration in different media and fermentation mutants. Related to Figure 4 and Figure 5.**

(A) The CV of the calibrated ATP concentration for the reference strain in different media. (B) The CV of the calibrated ATP concentration for the indicated genetic backgrounds in M9GlcCAA medium. Data are represented in boxplots, with the medians labeled by the circles and 25%-75% percentiles shown by the square boxes.

### Supplementary Methods

#### Estimate the ATP turnover rate per cell cycle

We calculated the estimated turnover rate of ATP per cell cycle based on measurements of ATP turnover rates per minute and growth rates of *Escherichia coli* cultures growing under different culture conditions[1].

| Media | growth rate[1] (1/hour) | ATP turnover rate[1] (1/min) | Calculated ATP turnover rate ($10^4$/cell cycle) |
|---|---|---|---|
| M9Glucose | 0.95 | 311 | 1.3 |
| M9Gluconate | 0.85 | 375 | 1.8 |
| M9Glycerol | 0.77 | 257 | 1.4 |
| M9Malate | 0.54 | 268 | 2.1 |
| M9Acetate | 0.36 | 453 | 5.2 |

#### Design and manufacture of the microfluidic device

The microfluidic design and setup (see figure below) were based on the "mother machine" and "chemoflux" devices[6,7].

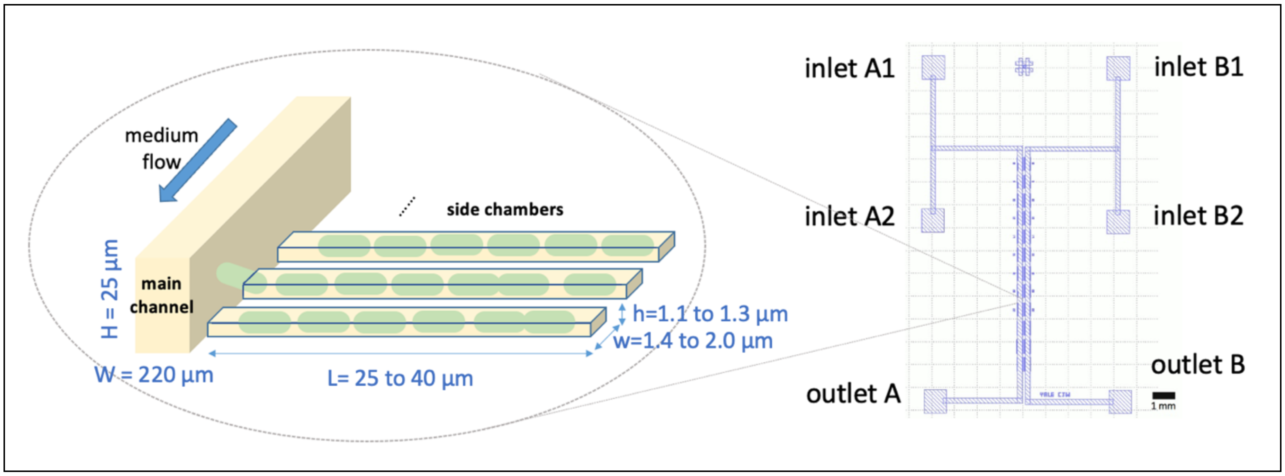

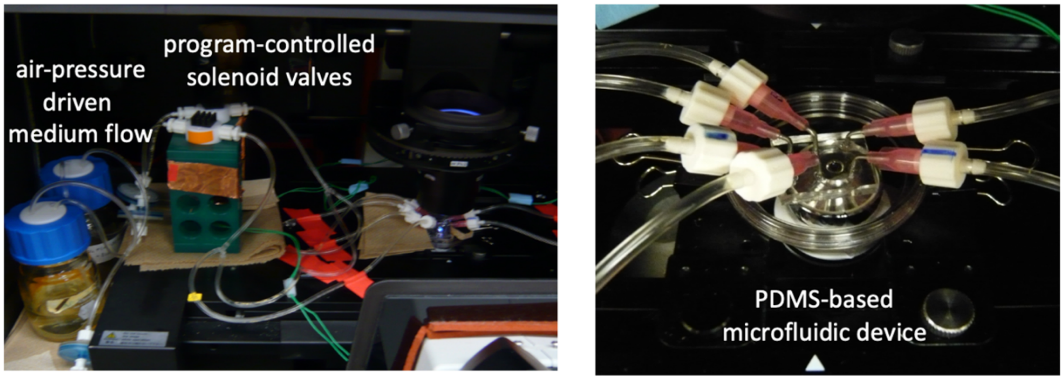


**Upper panel**: Two back-to-back microfluidic devices, each has two inlets and one outlet. The side chambers have a series of different width (w) and length (L), accomodating for different cell sizes in various nutrient conditions. Different height (h) of the side chamber can be achieved by varying the spinning speed of SU8 mold. **Lower panel: Photos of the** icrofluidic setup on the microscope with customized medium valves (left) and junctions (right).

We used the following protocol for manufacturing the SU8 master of microfluidic device:

**(1) Mask layout design**: The mask for the main channel and the side chambers were designed by KLayout software. The layout files were saved as GDS file and submitted to Yale SEAS Cleanroom kit. Both masks contain identical fiducial features on the edge for alignment purpose.

**(2) Mask generation**: This part was done by the SEAS Cleanroom staff, using Heidelberg DWL-66 Laser Mask Writer on 5 inches Chromium-coating glass masks.

**(3) Photolithography steps**:

(3-1) The 4 inches Silicon wafer (University Wafer) was first immersed in “Piranha” solution (sulfuric acid: hydrogen peroxide=3:1) for 10 min. After treatment, the wafer was rinsed with $H_2O$ 3 times and treated with the buffer oxide etch (BOE) (40% of $NH_4F$ with 50% of HF in 6:1 volume ratio) for about 1 min to create a hydrophobic surface.

(3-2) The wafer was baked on 95°C hot plate for 2 min to remove water.

(3-3) SU8 coating for 1st layer: A 1:1 mixture of photoresist SU8-2001 and SU-2002 were used for the first layer. For spreading the SU8 on the silicon wafer, following centrifugation protocol is used for generating a SU8 layer with thickness 1.0 to 1.2 µm.

**SU8 centrifugation procedure (first layer)**

STEP 1: 500 rpm for 10 sec

STEP 2: 3000 rpm for 45 sec

After spreading of SU8, the wafer was baked on a hotplate in the following procedure: (i) Put the wafer on the 65°C hotplate and raise the temperature of the hotplate from 65°C to 95°C with 0.5°C/sec speed. This takes 1 minute to reach 95°C. (ii) Bake the wafer in 95°C for 2 min. (iii) Leave the wafer on the hotplate and lower the temperature of the hotplate from 95°C to 65°C with 0.5°C/sec speed. (iv) Remove the wafer from hotplate and let it cool to room temperature.

(3-4) Photolithography for the 1st layer: The EVG 620 Mask aligner (EV Group) is used to perform the UV lithography step. Short wavelength UV was applied on the SU8 photoresist with exposure energy 3.75 mJ/cm$^2$, using 10 sec exposure time for three repeats. After exposure, the wafer was baked in 95°C for 2 min using the same procedure in step (3-3).

(3-5) SU8 coating for 2nd layer: The photoresist SU8-2010 was used to generate a thick SU8 layer around 30 µm. For spreading the SU8 on the silicon wafer, following centrifugation protocol is used:

**SU8 centrifugation procedure (second layer)**

STEP1: 500 rpm for 20 sec

STEP2: 2200 rpm for 45 sec

After spinning, the wafer with SU8 was baked in 95℃ for 2 min using the same procedure as in (3-3).

(3-6) Reveal the fiducial features of the first layer: To perform a proper alignment between first and second layer, the fiducial of first layer is revealed by dropping the Developing Solution (1-methoxy-2-propanol acetate) on the fiducial position until the pattern of the feature can be clearly seen. The wafer was dried by nitrogen gas.

(3-7) Photolithography for 2nd layer: Similar to step (3-4), the EVG 620 Mask aligner (EV Group) is used to perform the UV lithography. Short wavelength UV was applied on the SU8 photoresist with exposure energy 3.75 mJ/cm$^2$, using 10 sec exposure time for six repeats. After exposure, the wafer was baked in 95℃ for 6 min using the similar procedure in step (3-3).

(3-8) Full development of the wafer: After the photolithography step of the second layer, the wafer is fully developed by immersion of the wafer in the Developing Solution for 2-3 min. Same procedure is repeated until the edge of SU8 pattern can be clearly seen by eye. After the development step. The wafer is rinse with isopropanol for multiple times until the isopropanol solution does not become turbid of the wafer. The wafer is then dried with nitrogen gas.

(3-9) Measuring the height of SU8 features: the height of SU8 features, including the main trench and side chambers, by KLA-Tencor ASIQ profiler (KLA).

#### Luciferase assay and ATP level calibration

To generate a calibration curve between $R_{ATP}$ values (obtained by ratiometric imaging) and absolute ATP concentrations (measured using a luciferase assay), we varied the ATP level of *Escherichia coli* cells (strain CJW7556) by adding 0, 0.2 or 2 mM of potassium cyanide (KCN, final concentration) for 20 min. For ratiometric imaging, CJW7556 cells expressing QUEEN-2m were grown in a microfluidic device at 25°C first in M9GlcCAA until they reach exponential phase. Then, the culture medium was switched to M9GlcCAA containing 0, 0.2 or 2 mM KCN. The $R_{ATP}$

value obtained after 15 min of KCN treatment was used for calibration. For the luciferase assay, CJW7556 cells were grown in batch culture and treated with KCN under the same condition as for the ratiometric measurements. The luciferase assay was done using the BacTiter-Glo kit (Promega) following the manufacturer's instructions. ATP standards (ThermoFisher) were used in the luciferase assay to convert luminescence values into absolute number of ATP molecules.

To convert the number of ATP molecules into ATP concentration, the <u>total cell volume in the luciferase mixture</u> was estimated using by three numbers:

| Quantity | Number | Reference |
|---|---|---|
| Cell density per optical density (OD) per ml | $5.9 \times 10^8$ (cells/OD/ml) | [2] |
| Cell volume | 1.9 $\mu m^3$ | [3] |
| Fraction of cytoplasmic volume | 84% | [4] |

The above estimations allowed the conversion of the number of ATP molecules into a cytoplasmic ATP concentration. From the original report of the QUEEN-2m biosensor[5] the $R_{ATP}$ values are known to be linearly correlated with ATP concentration within a range of 1 to 5 mM. A linear fit (see following figure) was used to convert the $R_{ATP}$ data into ATP concentration data. The relation between $R_{ATP}$ value and ATP concentration is shown in Figure 1B.

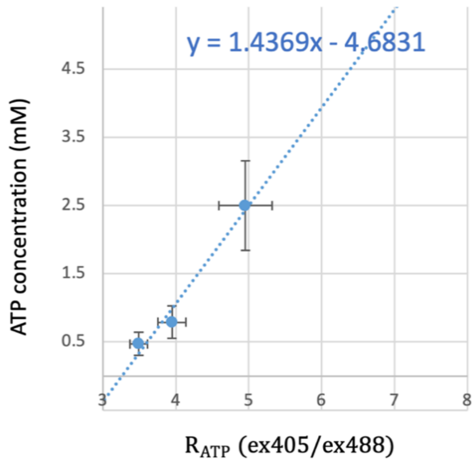


Calibration curve between ratiometric measurements ($R_{ATP}$) and absolute ATP concentrations (mM).

#### Determination of the biosensor protein level

To infer the QUEEN-2m protein level, the fluorescence signal of 405nm and 488nm excitation, denoted as $s_{405}$ and $s_{488}$, were used. As we shall see in below, the biosensor protein level is a weighted sum of $s_{405}$ and $s_{488}$. The excitation spectra of ATP-unbound QUEEN-2m and ATP-bounded QUEEN-2m is obtained from Fig 1e in the reference[5], where the 0 mM and 9.4 mM curves are used, respectively. Both bound and unbound states contribute to $s_{405}$ and $s_{488}$. The ATP-bound QUEEN-2m emits more signal under 405 nm excitation.

We used a linear combination of these signals to infer the biosensor protein level. To achieve this, we decomposed the biosensor into bounded and unbounded fractions, denoted as $n_b$ and $n_u$, where the total amount biosensor protein is equal to $n_b$+$n_u$. The signal under conditions (i) and (ii) is a linear combination of $n_b$ and $n_u$. Specifically, we can write

$$s_{405} = \phi_1(m_{11}u_b + m_{12}u_n)$$
$$s_{488} = \phi_2(m_{21}u_b + m_{22}u_n)$$

where the coefficient $m_{ij}$ can be obtained from the spectral property of biosensor protein, and the coefficient $\phi_k$ represents the amount of light source energy used for excitation. From the spectral measurement of the original paper (Figure 1e in reference [5]), we estimate the coefficient to be (in arbitrary unit):

$$M = \begin{pmatrix} m_{11} & m_{12} \\ m_{21} & m_{22} \end{pmatrix} \approx \begin{pmatrix} 7000 & 4900 \\ 1000 & 5200 \end{pmatrix}$$

The light source energy $\phi_1$ and $\phi_2$ is proportional to (LED power intensity) x (exposure time). Under our experimental setup, the LED used for 405 nm (SpectraX, Violet LED (395/25 nm)) and 488nm (SpectraX , Violet LED (470/24 nm)) has a power of 195 mW and 100 mW, respectively (see https://lumencor.com/products/spectra-x-light-engine/). In our experiments, the exposure times for 405nm and 488 nm were 200 ms and 400 ms, respectively. Hence, the coefficients of $\phi_k$ can be estimate as $\phi_1 = 195, \phi_2 = 200$ (arbitrary unit in energy). Given the above coefficients, the relation between signal and bounded/unbounded biosensor level has the following relation:

$$s_{405} = 195\,(7000\,u_b + 4900\,u_n)$$
$$s_{488} = 200\,(1000\,u_b + 5200\,u_n)$$

Solving for the equation, we have

$$u_b = 10^{-7}(8.47\,s_{405} - 7.78\,s_{488})$$
$$u_n = 10^{-7}(-1.63\,s_{405} + 11.11\,s_{488})$$

Therefore, the total amount of biosensor can be estimated as

$$u_b + u_n = 10^{-7}(6.84\,s_{405} + 3.33\,s_{488})$$

while the unit in above formula is arbitrary, the relative ratio (6.84 : 3.33) is important for inferring the relative amount of biosensor under the same experimental setup.
